# A Designed Ankyrin Repeat Protein (DARPin) Targeting EGFR Inhibits Ovulation and Enables a Novel Platform for Studying Ovarian Biology and Pathophysiology

**DOI:** 10.64898/2026.06.22.732379

**Authors:** Yubing Liu, Jiyang Zhang, Shichao Liu, Connie Mitra, Yue Liu, Hannah M. VanBenschoten, Brittany Goods, Feng Chen, Shuo Xiao

**Affiliations:** Department of Pharmacology and Toxicology, Ernest Mario School of Pharmacy, Rutgers University, Piscataway, NJ 08854, USA; Environmental and Occupational Health Sciences Institute (EOHSI), Rutgers University, Piscataway, NJ 08854, USA; Thayer School of Engineering, Dartmouth College, Hanover NH 03755, USA; School of BioSciences, The University of Melbourne, Parkville VIC 3052, Australia; Department of Radiology, Weill Cornell Medicine, New York, NY 10021, USA

**Keywords:** DARPin, EGFR, ovulation, ovarian follicle, ovarian biology, *ex vivo* follicle culture

## Abstract

Ovarian disorders, including anovulation, primary ovarian insufficiency (POI), and polyendocrine metabolic ovarian syndrome (PMOS), affect millions of reproductive-age women worldwide; however, mechanistic studies of ovarian biology and pathophysiology remain challenging because current experimental approaches often lack selectivity, tunability, or physiological relevance. Genetically modified animal models are labor-intensive and irreversible; small molecules frequently exhibit off-target effects; and conventional antibodies have limited tissue penetration and restricted temporal control. Designed ankyrin repeat proteins (DARPins) represent a highly modular protein engineering platform with advantages in specificity, size, stability, and extracellular targeting, but their utility in reproductive biology remains largely unexplored. Here, we used epidermal growth factor receptor (EGFR)-targeting DARPins as a proof-of-concept platform to interrogate ovarian signaling during ovulation. Screening of engineered anti-EGFR DARPins identified SX-006, a bispecific tetravalent construct with robust cross-species EGFR binding and potent biological activity. Using an *ex vivo* murine ovulation system, SX-006 inhibited follicle rupture in a dose-dependent manner with IC50 of 1.21 μM without overt cytotoxicity. Lower concentrations of SX-006 preferentially perturbed follicle rupture while largely preserving oocyte meiotic maturation and luteinization, suggesting differential sensitivity of ovulatory processes to extracellular EGFR blockade. Comparative transcriptomic analyses further revealed that extracellular EGFR blockade and small molecule-based intracellular EGFR kinase inhibition produce overlapping but also distinct transcriptional responses, supporting biologically distinct modes of ovulatory signaling pathway perturbation. Together, these findings establish DARPins as a selective, tunable, and physiologically relevant platform for studying ovarian signaling and provide proof-of-concept for extracellular receptor targeting in ovarian biology, infertility research, and non-hormonal contraceptive development.

**Summary sentence:** An engineered EGFR-targeting DARPin selectively inhibits ovulation through extracellular receptor blockade and establishes a versatile platform for investigating ovarian signaling and reproductive disorders.

## Introduction

Up to 15-20% of women of reproductive age experience infertility, with ovarian disorders as a leading cause, including primary ovarian insufficiency (POI), polycystic ovary syndrome (PCOS, now termed polyendocrine metabolic ovarian syndrome or PMOS), anovulation, and aneuploidy[1–3]. Despite its importance, mechanistic studies of ovarian biology and pathophysiology remain challenging due to the lack of precise and scalable approaches. Genetically modified animal models have provided critical insights; however, the process is time-consuming, costly, and not readily adaptable for reversible or dose-dependent interrogation of targeted signaling pathways. Small molecules are often limited by pharmacokinetic constraints, including rapid clearance and variable bioavailability [4]. Small molecules also often exhibit off-target effects, complicating the interpretation of complex signaling networks governing biological functions [5, 6]. The relatively large antibodies exhibit high specificity targeting ligand–receptor interactions. However, they restrict tissue penetration within complex ovarian structures. Moreover, the Fc-mediated effector function and long systemic half-life may introduce confounding biological effects and limit temporal control of targeted signaling [7]. The antibody engineering and production can also be costly and less amenable to rapid iteration. Collectively, these limitations underscore a critical need for new tools that enable selective, tunable, and physiologically relevant modulation of ovarian signaling.

Designed Ankyrin Repeat Proteins (DARPins) emerge as a precision targeting tool to address the limitations of both small molecules and monoclonal antibodies [8, 9]. DARPins are highly stable, engineered scaffold proteins derived from naturally occurring ankyrin repeat motifs [10]. Unlike small molecules, which often exhibit off-target effects, DARPins enable highly selective extracellular targeting of defined receptor epitopes, thereby enabling pathway-specific modulation of ligand-receptor interactions [8]. Compared with traditional monoclonal antibodies, DARPins lack disulfide bonds and Fc regions, allowing for cost-effective, high-yield expression in *Escherichia coli*, while minimizing undesirable immune effector functions [9, 11]. The compact molecular size (∼14–18 kDa per domain) of DARPins provides a rigid framework for high-affinity molecular recognition and enables facile engineering into multi-specific or multivalent formats [10]. Moreover, bispecific or biparatopic DARPins engineered with a leucine zipper can also homodimerize to form tetravalent complexes that can cross-link surface receptors [12, 13]. This architectural advantage enables specific DARPin constructs to potently inhibit receptor recycling and down-regulate surface receptors [12]. Because of their robust biophysical properties and potent antagonistic capabilities, DARPins have emerged as an ideal tool for interrogating coordinated ligand-receptor systems.

Ovulation is an essential reproductive event in which a fully grown preovulatory follicle releases a mature oocyte or egg in response to the pituitary luteinizing hormone (LH) surge. This tightly controlled process sustains female reproductive cycles and fertility and serves as a hormonal and non-hormonal contraceptive target [14, 15]. Impaired ovulation is a hallmark of PCOS and luteinized unruptured follicle (LUF) syndrome, two prevalent ovarian disorders affecting adolescents and young adult women [16, 17]. The epidermal growth factor receptor (EGFR) plays a central role in the ovarian response to the LH surge. The binding of LH to its membrane receptor LHCGR induces EGF-like ligands in granulosa cells, including amphiregulin (AREG), epiregulin (EREG), and betacellulin (BTC)[18–20]. These EGF ligands bind to EGFR and activate extracellular signal-regulated kinase (ERK) signaling cascades, which are essential for cumulus expansion, oocyte meiotic resumption, and follicle rupture. So far, most studies investigating EGFR signaling rely on genetically modified mouse models or small-molecule tyrosine kinase inhibitors [21–23]. While these models disrupt EGFR expression or intracellular kinase activity, they do not target ligand-dependent EGFR activation and often broadly affect downstream signaling pathways. Moreover, EGFR ligands bind to the extracellular domain of EGFR; thus, experimental tools that selectively block extracellular ligand-EGFR interactions may provide a precise strategy to elucidate EGF ligand-EGFR-mediated ovulatory signaling.

In the present study, we used anti-EGFR DARPins in combination with a 3D *in vitro* ovarian follicle growth (IVFG) and *ex vivo* ovulation system as a novel platform to study ovarian signaling during ovulation. We demonstrate that by selectively targeting the extracellular ligand-receptor interactions, anti-EGFR DARPin inhibits ovulation in a dose-dependent manner without overt cytotoxicity and enables precise modulation of EGFR signaling distinct from a small-molecule EGFR inhibitor that suppresses intracellular EGFR kinase activity. Our findings establish DARPin as a versatile and tunable platform for studying ovarian signaling and highlight its therapeutic potential for ovarian disorders, infertility, and the development of non-hormonal contraceptives.

## MATERIALS AND METHODS

### Cloning, Expression, and Purification of anti-EGFR DARPins

The design and recombinant expression strategy for anti-EGFR DARPins were adapted from previously established protocols [13]. The genes encoding the anti-EGFR DARPins were optimized for E. coli expression and cloned into the pET28a(+) expression vector under the control of a T7 promoter. Each construct was designed with an N-terminal 6xHis-tag followed by a tobacco etch virus (TEV) protease cleavage site to facilitate affinity purification. Plasmids were transformed into E. coli BL21(DE3) competent cells. Bacterial cultures were grown in Luria-Bertani (LB) medium supplemented with 50 µg/mL kanamycin at 37°C until the optical density at 600 nm reached 0.6-0.8. Recombinant protein expression was induced via the addition of 0.5 mM isopropyl β-D-1-thiogalactopyranoside (IPTG), followed by an incubation at 37°C for 3-4 h. Cells were harvested by centrifugation and resuspended in a lysis buffer (20 mM Sodium Phosphate, pH 7.4, 500 mM NaCl, 10 mM imidazole) containing 0.5 mg/mL lysozyme and an EDTA-free Protease Inhibitor Cocktail. Lysis was achieved by sonication on ice, and soluble proteins were recovered from the supernatant after centrifugation. The DARPins were isolated using nickel-nitrilotriacetic acid (Ni-NTA) affinity chromatography; the column was washed with buffer containing 20 mM imidazole, and the target proteins were eluted with 500 mM imidazole. Eluted fractions were dialyzed against 1x PBS (pH 7.4) and passed through a 0.22 µm filter.

### SDS-PAGE Analysis

Protein purity and anticipated molecular weights were verified by sodium dodecyl sulfate-polyacrylamide gel electrophoresis (SDS-PAGE). DARPin samples were diluted in 2x Laemmli buffer, heated at 95°C for 5 minutes, and loaded onto 8–16% polyacrylamide gels (BioRad, Cat. No. 456–1105). Electrophoresis was carried out at 120 V for 45 minutes, after which the gels were stained using Coomassie Brilliant Blue to visualize the protein bands. Gel images were acquired using a Bio-Rad imaging system. Band intensities were quantified using ImageJ software (NIH, Bethesda, MD).

### Endotoxin Removal of Anti-EGFR DARPins

To prevent endotoxin interference in downstream biological assays, the purified DARPins were processed using Pierce High-Capacity Endotoxin Removal Spin Columns (Thermo Scientific, Cat. No. 88274) equilibrated with endotoxin-free water. Following a 1-hour incubation at room temperature to facilitate binding, samples were centrifuged at 500 g for 2 minutes to recover the endotoxin-depleted proteins. Final endotoxin levels were confirmed to be below the assay detection limit using the Pierce Rapid Gel Clot Endotoxin Assay Kit (0.5 EU/mL sensitivity, Cat. No. A43882). Protein concentration was measured using a NanoDrop spectrophotometer by absorbance at 280 nm. The concentration of each purified DARPin was calculated based on its theoretical molecular weight and extinction coefficient. Total protein yield was calculated by multiplying the measured protein concentration by the final recovered volume. Purified proteins were aliquoted and stored at 4°C until use.

### Enzyme-Linked Immunosorbent Assay (ELISA)

Binding affinities and cross-reactivity were assessed using a quantitative ELISA. High-binding 96-well microplates were coated with either human (Sino Biological, Cat. No. 10001-H02H) or mouse (Sino Biological, Cat. No. 51091-M02H) recombinant EGFR protein and incubated overnight. The wells were subsequently blocked with 1% BSA in PBS for 1 hour at room temperature. Serial dilutions of the DARPins were added to the wells and incubated for 1 hour. Following washing steps, specific DARPin binding was detected using a biotinylated anti-HisTag monoclonal antibody, followed by a 30-minute incubation with streptavidin-horseradish peroxidase (HRP). The colorimetric reaction was developed utilizing 3,3’,5,5’-tetramethylbenzidine (TMB) substrate and terminated with an ELISA stop solution. Absorbance was recorded at 450 nm using a microplate reader.

### Animals

CD-1 mice were used for ovarian follicle isolation. Mice were obtained from an existing breeding colony originally purchased from Envigo and maintained at Rutgers University. Animals were housed in polypropylene cages in the Animal Care Facility of the Research Tower at Rutgers University under controlled conditions of temperature (22 ± 1°C) and humidity (30%–70%), with a 12 h light/12 h dark cycle. All animals were maintained and handled in accordance with the National Institutes of Health Guide for the Care and Use of Laboratory Animals and protocols approved by the Rutgers University Institutional Animal Care and Use Committee (IACUC).

### Assessment of anti-EGFR DARPin effects in the eIVFG-based in vitro ovulation model

To evaluate the effects of anti-EGFR DARPin on hCG-induced ovulatory signaling, immature preantral follicles were isolated from 16-day-old CD-1 female mice and cultured using our previously established three-dimensional encapsulated in vitro follicle growth system. Briefly, ovaries were collected after CO_₂_ euthanasia and incubated in L15 medium (Invitrogen) containing Liberase (Sigma-Aldrich) and DNase I for enzymatic follicle release. Secondary follicles with diameters of approximately 150–180 μm were manually isolated under a stereomicroscope and encapsulated individually in 0.5% alginate hydrogel (Sigma-Aldrich). Encapsulated follicles were cultured individually in 96-well plates in follicle growth medium supplemented with 10 mIU/mL recombinant FSH (rFSH; Organon, gifted by Dr.Mary Zelinski from the Oregon Nonhuman Primate Research Center at the Oregon Health and Science University, Beaverton, Oregon, USA),. Follicles were maintained in eIVFG culture until the preovulatory stage. To induce ovulation, follicles were stimulated with 1.5 IU/mL hCG (Sigma-Aldrich) and 10 mIU/mL rFSH. At the time of hCG stimulation, follicles were simultaneously treated with vehicle control, Tyrphostin AG1478 (Selleckchem, Cat. No. S2728), a selective EGFR tyrosine kinase inhibitor, at 1 μM as a positive EGFR inhibition control, anti-HSA DARPin at 10 μM as a negative DARPin control, or anti-EGFR DARPins. The anti-HSA DARPin was included to control for nonspecific effects of the DARPin scaffold.

Follicles and conditioned media were collected at defined time points after hCG stimulation for downstream analyses. Follicles collected at 4 h after hCG stimulation were used for qRT-PCR analysis of ovulation-related gene expression and RNA-seq analysis. Ovulation was assessed at 16 h after hCG treatment using an Olympus inverted microscope equipped with a 10× objective (Olympus Optical Co. Ltd., Tokyo, Japan) and TCapture imaging software (Tucsen, v5.1.1). A follicle was defined as “ruptured” when one side of the follicular wall was breached, whereas follicles with an intact follicular wall were classified as “unruptured.” For IC50 estimation, follicle rupture inhibition was calculated by normalizing each treatment group to the vehicle control within the same independent experiment. Inhibition values were fitted using nonlinear regression with a four-parameter logistic dose–response model in GraphPad Prism. Because the data were normalized to percent inhibition, the bottom and top of the curve were constrained to 0% and 100%, respectively. Oocytes from all follicles included in the rupture assay were examined for extrusion of the first polar body. Oocytes with a visible first polar body were classified as polar body extrusion oocytes. Oocytes that remained at the germinal vesicle (GV) stage or underwent germinal vesicle breakdown (GVBD) without first polar body extrusion were classified as non-polar body extrusion oocytes. Following ovulation induction, follicles were continuously cultured in ovulation induction medium without rFSH for 48 h to induce luteinization and progesterone secretion. Conditioned media were collected at 48 h and stored at −20°C until progesterone measurement by ELISA.

### Follicular somatic cell number and viability assay

Follicular somatic cell number and viability were assessed at 16 h after hCG treatment. Follicles were collected and dissociated into single cells using Accutase (Innovative Cell Technologies, Cat. No. AT104). The resulting cell suspension was mixed with 0.4% trypan blue, and viable and non-viable cells were counted using a hemocytometer. Unstained cells were considered viable, whereas trypan blue-positive cells were considered non-viable. Total follicular somatic cell number and cell viability were calculated for each follicle. A total of 5–6 follicles were analyzed per treatment group.

### Progesterone (P4) ELISA assay

P4 concentrations in the conditioned media were measured using a progesterone ELISA kit (Cayman Chemical, Ann Arbor, MI; RRID: AB_2811273; Cat. No. 582601) according to the manufacturer’s instructions. Briefly, mouse anti-rabbit IgG-precoated wells were incubated with standards, conditioned culture media, rabbit antiserum, and progesterone-acetylcholinesterase (AChE) conjugate for 60–120 min. After incubation, the wells were washed three times with washing buffer, then Ellman’s reagent was added and incubation continued at room temperature for 60–90 min. Absorbance was measured at 414 nm using a BioTek SpectraMax M3 microplate reader (BioTek Instruments, Inc., Winooski, VT) within 15 min. The reportable range of the P4 assay was 7.8–1,000 pg/mL. The assay sensitivity, defined by the manufacturer as the analyte concentration corresponding to 80% B/B0 (% Bound/Maximum Bound) and derived from the standard curve provided in the kit manual, was 10 pg/mL. The reported inter-assay coefficient of variation (CV) for P4 was 7.7%–16.4%, and the intra-assay CV was 7.3%–54.5%. Conditioned media from 9–10 follicles per treatment group were used for P4 measurement.

### RNA extraction and RT-qPCR

To examine the effects of anti-EGFR DARPin on ovulatory gene expression, follicles were collected at 4 h after hCG stimulation for RT-qPCR analysis, because many LH/hCG-responsive genes are strongly induced at this time point in both in vivo and in vitro ovulation models. Total RNA was extracted from individual follicles using the PicoPure RNA Isolation Kit according to the manufacturer’s instructions. RNA concentration and purity were assessed using a NanoDrop spectrophotometer. Total RNA was reverse-transcribed into cDNA using the SuperScript III First-Strand Synthesis System with random hexamer primers (Invitrogen, Cat. No. 18080400). RT-qPCR was performed in 384-well plates using Power SYBR Green PCR Master Mix on a ViiA 7 Real-Time PCR System. The cycling conditions were 95°C for 10 min, followed by 40 cycles of 95°C for 15 s and 60°C for 40 s. A melting curve analysis was performed to confirm primer specificity. The expression of ovulation-related genes, including *Pgr*, *Ptgs2*, *Has2*, *Ereg*, *Sult1e1,* and *Star,* was examined. Relative gene expression was normalized to *Hprt*. Primer sequences are listed in Table S2. A total of n = 5 or 6 follicles per treatment group were analyzed.

### Single-follicle RNA-sequencing (RNA-seq) analysis

To evaluate the effect of anti-EGFR DARPin on hCG-induced follicular transcriptomic responses, follicles were collected at 4 h after hCG stimulation for single-follicle RNA-seq analysis. Follicles were treated with vehicle control or anti-EGFR DARPin SX-006 at the time of hCG-induced ovulation. A total of n = 4 follicles per group were used for RNA-seq analysis. Total RNA was extracted from individual follicles using the Arcturus PicoPure RNA Isolation Kit according to the manufacturer’s instructions. Library preparation and low-input RNA-seq were performed on the Illumina NovaSeq X Plus PE150 platform by Novogene Corporation. High-quality trimmed paired-end reads were processed using Galaxy (version 24.2.rc1). RNA-seq reads were aligned to the whole mouse genome assembly-mm10 by HISAT2, quantified by featureCounts, and normalized using the Transcripts Per Million (TPM) method.

Principal component analysis was performed in R (version 4.3.3) to assess sample clustering. Differential gene expression analysis was conducted using DESeq2 by comparing SX-006-treated follicles with vehicle-treated control follicles. Differentially expressed genes were defined as genes with a fold change ≥ 2.0 or ≤ 0.5 and an FDR-adjusted p-value < 0.05. Gene Ontology and pathway enrichment analyses were performed using Bioconductor packages in R, and enrichment plots were generated using ggplot2.

To compare the transcriptional effects of SX-006 with pharmacological EGFR kinase inhibition, DEGs identified in SX-006-treated follicles were compared with DEGs from AG1478-treated follicles collected 4 h after hCG stimulation using the same in vitro ovulation system. The AG1478 RNA-seq data were generated from an independent experimental batch in our laboratory under the same experimental and sequencing procedures as the other samples. Shared and treatment-specific DEGs were identified, followed by functional enrichment analyses to define common and distinct biological processes affected by SX-006-mediated extracellular EGFR blockade and AG1478-mediated intracellular kinase inhibition.

### RNA-seq data availability

All raw FASTQ files and raw count data from single-follicle RNA-seq experiments have been deposited in the Gene Expression Omnibus (GEO) database under accession numbers GSE334906 and GSE335098.

### Literature-curated EGFR-associated module construction and GSEA analysis

To facilitate pathway-level comparison of extracellular EGFR blockade and intracellular EGFR kinase inhibition, we curated 12 EGFR-associated gene modules based on established ovulatory signaling pathways and EGFR biology established in previous studies (Figure S4) [19, 24–36]. Modules were organized into four biological categories: (1) Core ovulatory EGFR programs (Modules 1–7); (2) Early luteal transition program (Module 8), (3) Canonical EGFR regulatory programs (Modules 9–11); and (4) oocyte-associated program (Module 12). Gene selection was based on published studies of LH/EGFR signaling, ovulation, cumulus expansion, prostaglandin biosynthesis, follicle rupture, luteinization, EGFR feedback regulation, receptor internalization, and oocyte meiotic control.

Gene set enrichment analysis (GSEA) was performed using GSEAPreranked (Broad Institute). Genes from each RNA-seq comparison were ranked according to the DESeq2 Wald statistic (“stat” value). Custom gene sets were formatted as GMT files and analyzed using 1,000 phenotype permutations with weighted enrichment statistics. Normalized enrichment scores (NES) and false discovery rate (FDR) q values were calculated for each module. Consistent with GSEA recommendations, modules with FDR q values < 0.25 were considered significantly enriched. Positive NES values indicate relative enrichment of a module, whereas negative NES values indicate relative suppression of the corresponding biological program.

### Data and Statistical Analysis

Data were analyzed using GraphPad Prism and R. For follicle rupture and oocyte meiotic maturation assays, follicle-level outcomes were first summarized within each independent experiment, and each independent experiment was treated as one biological replicate for statistical analysis. Follicle rupture rate was calculated as the percentage of ruptured follicles among the total follicles examined. Polar body extrusion rate was calculated as the percentage of oocytes with first polar body extrusion among all oocytes assessed.

For dose–response analysis, follicle rupture inhibition was normalized to the vehicle control within the same independent experiment using the following formula: inhibition (%) = [(vehicle rupture rate − treatment rupture rate) / vehicle rupture rate] × 100. IC50 values were estimated by nonlinear regression using a four-parameter dose–response inhibition model with variable slope. The bottom and top were constrained to 0% and 100%, respectively.

For continuous outcomes, including progesterone concentration, granulosa cell number, cell viability, and RT-qPCR gene expression, data are presented as mean ± SEM unless otherwise stated. RT-qPCR data were normalized to *Hprt*, and relative gene expression was calculated using the ΔΔCt method. Comparisons among multiple treatment groups were performed using one-way ANOVA followed by Dunnett’s multiple-comparisons test when each treatment was compared with the vehicle control. For comparisons between two groups, an unpaired two-tailed Student’s t-test was used. A P value < 0.05 was considered statistically significant.

For RNA-seq analysis, differential gene expression was analyzed using DESeq2. Differentially expressed genes were defined as genes with fold change ≥ 2 or ≤ 0.5 and FDR-adjusted *P* value < 0.05. Gene Ontology and KEGG pathway enrichment analyses were performed using R/Bioconductor packages.

## RESULTS

### Design and expression of various anti-EGFR DARPins and control constructs

To develop a precise tool for studying physiologically relevant extracellular EGFR signaling, we designed a comprehensive panel of anti-EGFR DARPins. The amino acid sequences for target-specific constructs were first adapted, with minor modifications, from previously published anti-EGFR DARPins [12]. The panel features monovalent DARPins targeting distinct EGFR epitopes, including SX-002 derived from the anti-EGFR_E01 sequence and SX-003 from the anti-EGFR_E69 sequence (Figure S1). To evaluate whether increased valency and altered spatial geometry could enhance receptor antagonism, we further engineered two bispecific constructs linking the E69 and E01 domains. These include SX-005 which used a flexible (G_4_S)_3_ linker adapted from the E69-GS-E01 design (Figure S1) and SX-006 that incorporates a central leucine zipper motif adapted from E69-LZ3-E01 to drive homodimerization (Figure 1A). As shown in Figure 1B, the bivalent single chain of SX-006 (E69-LZ3-E01) construct could undergo homodimerization mediated by the internal leucine zipper motif and assemble into a functional tetravalent binding complex capable of enhanced receptor cross-linking [12]. A non-targeting anti-human serum albumin (HSA) DARPin, SX-007 (Figure S1), derived from anti-HSA_v01 [37], was also expressed as a negative control.

**Figure 1.**
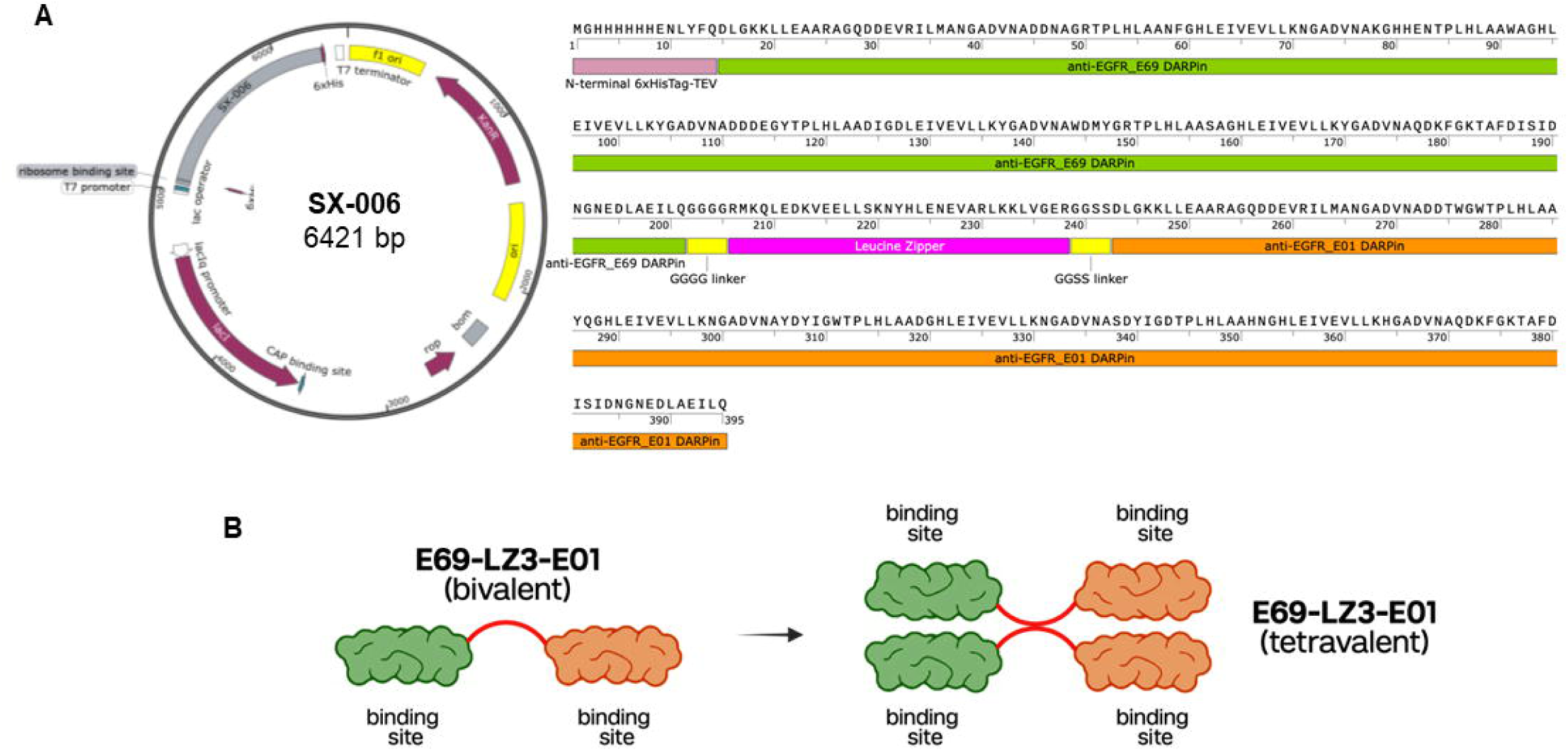
Design and architecture of the bivalent anti-EGFR DARPin SX-006 construct. (A) Plasmid map and corresponding amino acid sequence of the SX-006 (anti-EGFR bispecific DARPin_E69-LZ3-E01) expression vector (6421 bp). The modular pET28a(+) construct features a T7 promoter, an N-terminal 6xHis-tag, and a TEV protease cleavage site, followed sequentially by the anti-EGFR E69 DARPin domain, a leucine zipper motif, and the anti-EGFR E01 DARPin domain. (B) Schematic representation of the structural valency of the SX-006 (E69-LZ3-E01) construct. The bivalent single chain homodimerizes via its internal leucine zipper motif, forming a functional tetravalent binding complex capable of enhanced receptor cross-linking.

All DARPin constructs were cloned into a pET28a(+) expression vector, featuring an N-terminal 6xHis-tag and a TEV protease cleavage site to facilitate standardized affinity purification [13]. Recombinant expression was carried out in *E. coli* BL21(DE3). As summarized in Table S1, the bacterial expression system yielded robust quantities of soluble DARPin proteins, averaging 50.7-108.9 mg/L across batches. Following nickel-nitrilotriacetic acid (Ni-NTA) affinity chromatography, the purity and anticipated molecular weights of the constructs were validated via SDS-PAGE (Figure S2). These results confirm the successful production of high-purity DARPin complexes suitable for downstream functional assays.

### Cross-species analysis identifies SX-006 as an optimal DARPin for functional assays in the murine *ex vivo* follicle culture and ovulation assay

Because our *ex vivo* 3D ovarian follicle culture and ovulation model is murine-derived [15, 38–51], it was imperative to identify a DARPin construct capable of cross-reactivity with murine EGFR. We systematically evaluated the binding avidities of the purified DARPin panel against human and mouse recombinant EGFR using ELISA kits (Figure 2A). When tested against human EGFR, all anti-EGFR DARPins showed concentration-dependent binding but with distinct affinities. The monovalent SX-002 and bispecific SX-006 constructs exhibited a strong binding affinity with EC_50_ of 39.2 nM and 59.1 nM, respectively (Figure 2B). There were marked differences when evaluating binding avidities to murine EGFR. While SX-002 bound to human EGFR tightly, it was a weak binder of murine EGFR (Figure 2C). Similarly, the flexible-linker bispecific construct SX-005 showed weak binding to mouse EGFR (Figure 2C). The bispecific leucine-zipper construct SX-006 was the best binder against murine EGFR among all tested DARPins, maintaining an EC_50_ of 39 µM (Figure 2C). The non-targeting anti-HSA DARPin SX-007 and the PBS negative control exhibited partial to no specific binding to both human and mouse EGFR (Figure 2B-2C).

**Figure 2.**
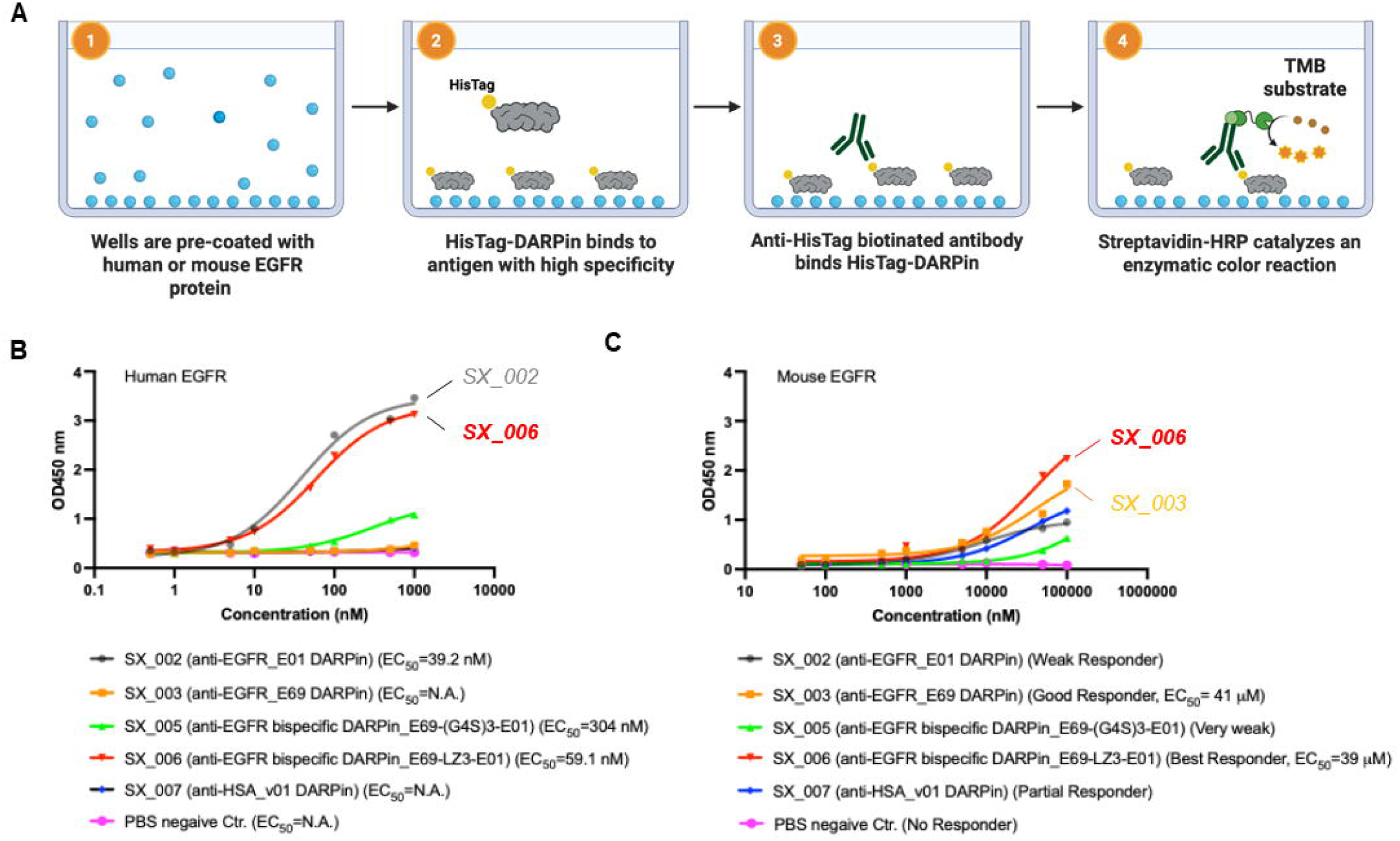
Evaluation of human and mouse EGFR cross-reactivity among the DARPin panel via ELISA. (A) Schematic workflow of the colorimetric anti-EGFR DARPin ELISA. Recombinant human or mouse EGFR is immobilized on microplate wells (1), followed by the specific binding of His-tagged DARPins (2). Detection is achieved using a biotinylated anti-HisTag antibody (3) and a subsequent streptavidin-HRP enzymatic reaction utilizing TMB substrate (4). (B) Concentration-dependent binding curves demonstrating high-affinity interactions with human EGFR. (C) Concentration-dependent binding curves against mouse EGFR. The bispecific DARPin SX-006 exhibits the most robust cross-species binding profile, justifying its selection for murine biological assays

### Effects of multiple anti-EGFR-DARPins on *ex vivo* follicle ovulation

SX-002 and SX-006 anti-EGFR DARPins were selected for functional assays using our *ex vivo* ovulation assay, with AG1478, a small molecule EGFR inhibitor, as a positive control, and anti-HSA DARPin as a negative control. Follicles treated with anti-HSA DARPin exhibited comparable outcomes of all examined endpoints to the vehicle group, including follicle rupture, oocyte polar body extrusion, progesterone secretion, follicular somatic cell numbers, and cell viability (Figure 3A-3H). Both SX-002 and SX-006 anti-EGFR DARPins concentration-dependently inhibited follicle rupture, with IC50 of 5.33 μM and 1.21 μM, respectively (Figure 3A-3C). SX-002 at all concentrations and SX-006 at 1 and 3 μM did not affect oocyte polar body extrusion; however, there were only 14.1 ± 7.1% oocytes from follicles treated with 10 μM SX-006 extruded the first polar body (Figure 3D-3E).

**Figure 3.**
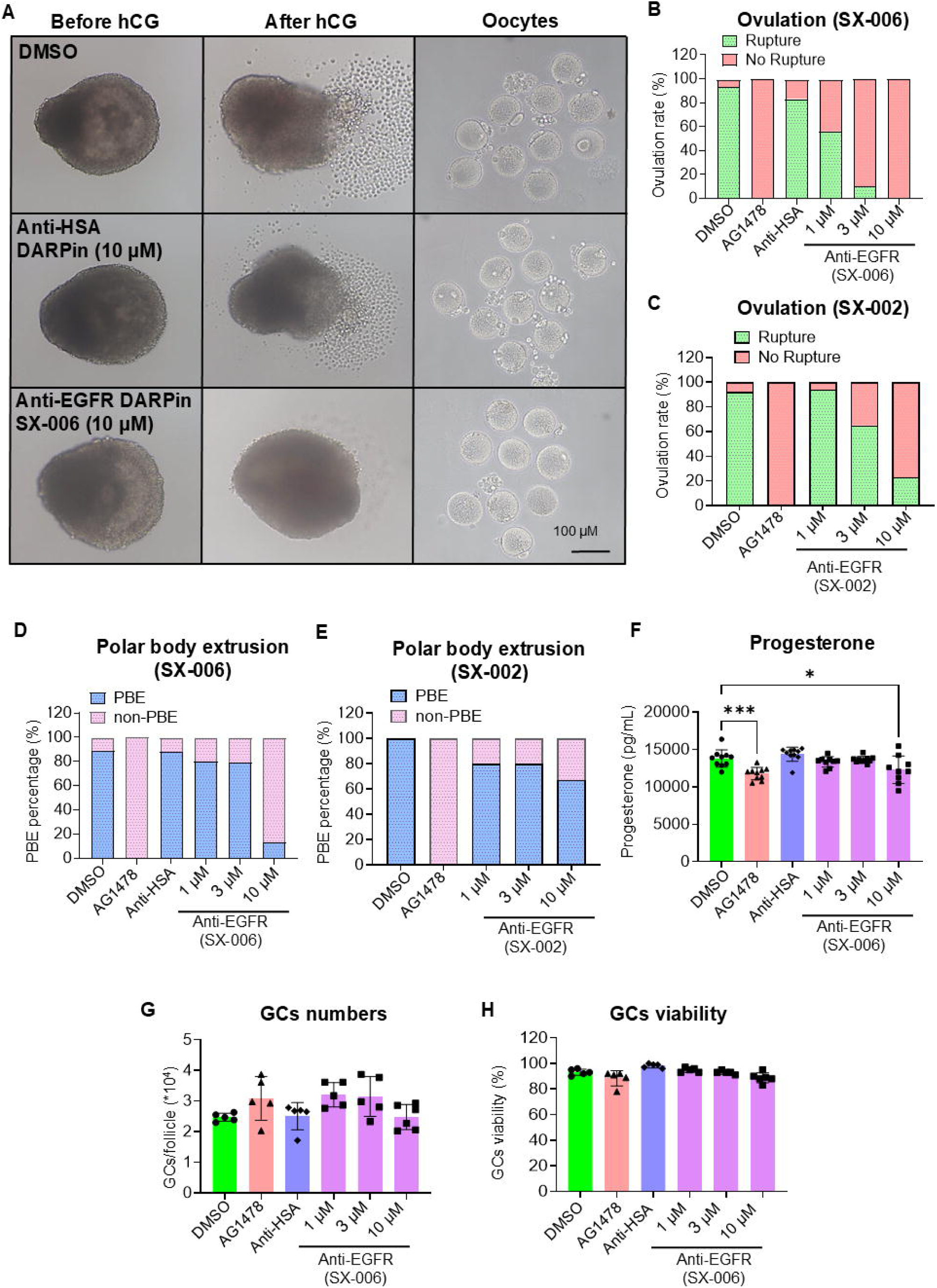
Anti-EGFR DARPins inhibit hCG-induced ovulation in the eIVFG model. (A) Representative bright-field images of follicles and oocytes after hCG-induced ovulation. DMSO and anti-HSA DARPin controls showed follicle rupture, whereas 10 μM SX-006 reduced rupture. Scale bar, 100 μm. (B, C) Follicle rupture rates after treatment with DMSO, Tyrphostin AG1478, anti-HSA DARPin, SX-006, or SX-002. SX-006 and SX-002 were tested at graded concentrations of 1, 3, and 10 μM. (D, E) Oocyte meiotic maturation, assessed by first polar body extrusion, following SX-006 or SX-002 treatment. (F) Progesterone levels in conditioned media after ovulation induction and luteinization culture. (G, H) Granulosa cell number and viability after treatment (n = 5). Data are presented as mean ± SEM. Unless otherwise indicated, n = 8–10 follicles per treatment group. Statistical significance was determined using one-way ANOVA followed by Dunnett’s multiple comparisons test. Asterisks indicate significant differences compared with the vehicle control group. \**P* < 0.05, \*\**P* < 0.01, and \*\*\**P* < 0.001.

The remaining follicular somatic cells after ovulation were cultured with hCG for 2 days to allow for luteinization and progesterone secretion. SX-006 with higher EGFR binning and greater ovulation-inhibition potency was selected for examining its effects on follicular cell survival and steroidogenic activity. SX-006 at lower concentrations of 1 μM and 3 μM did not alter progesterone secretion of the formed CL spheroids; however, similar to the positive control AG1478, SX-006 at 10 μM significantly reduced progesterone secretion (Figure 3F). The analysis of follicular somatic cell numbers and viability revealed no significant differences between all treatment groups, suggesting that the inhibited follicle rupture, oocyte meiotic resumption, and progesterone secretion were not attributable to DARPin-induced cytotoxicity (Figure 3G-3H). Together, these results indicate that anti-EGFR DARPins, particularly SX-006, selectively inhibit ovulation.

### Effects of anti-EGFR DARPin SX-006 on ovulatory marker genes

To identify molecular changes of SX-006-treated follicles, we next conducted a similar exposure experiment and collected follicles at 4 h post-hCG treatment during *ex vivo* ovulation induction to examine the expression of several established ovulatory genes using single-follicle RT-qPCR. The names, functions, and references of these genes are summarized in Table S3. Follicles exposed to anti-HSA DARPin showed expression levels comparable to those of vehicle-treated controls for all examined genes (Figure 4). However, anti-EGFR DARPin SX-006 significantly attenuated the hCG-induced upregulation of ovulatory genes, including *Pgr*, *Ptgs2*, *Has2*, *Ereg*, *Sult1e1*, and *Star*, with the most pronounced effects observed at 10 μM (Figure 4).

**Figure 4.**
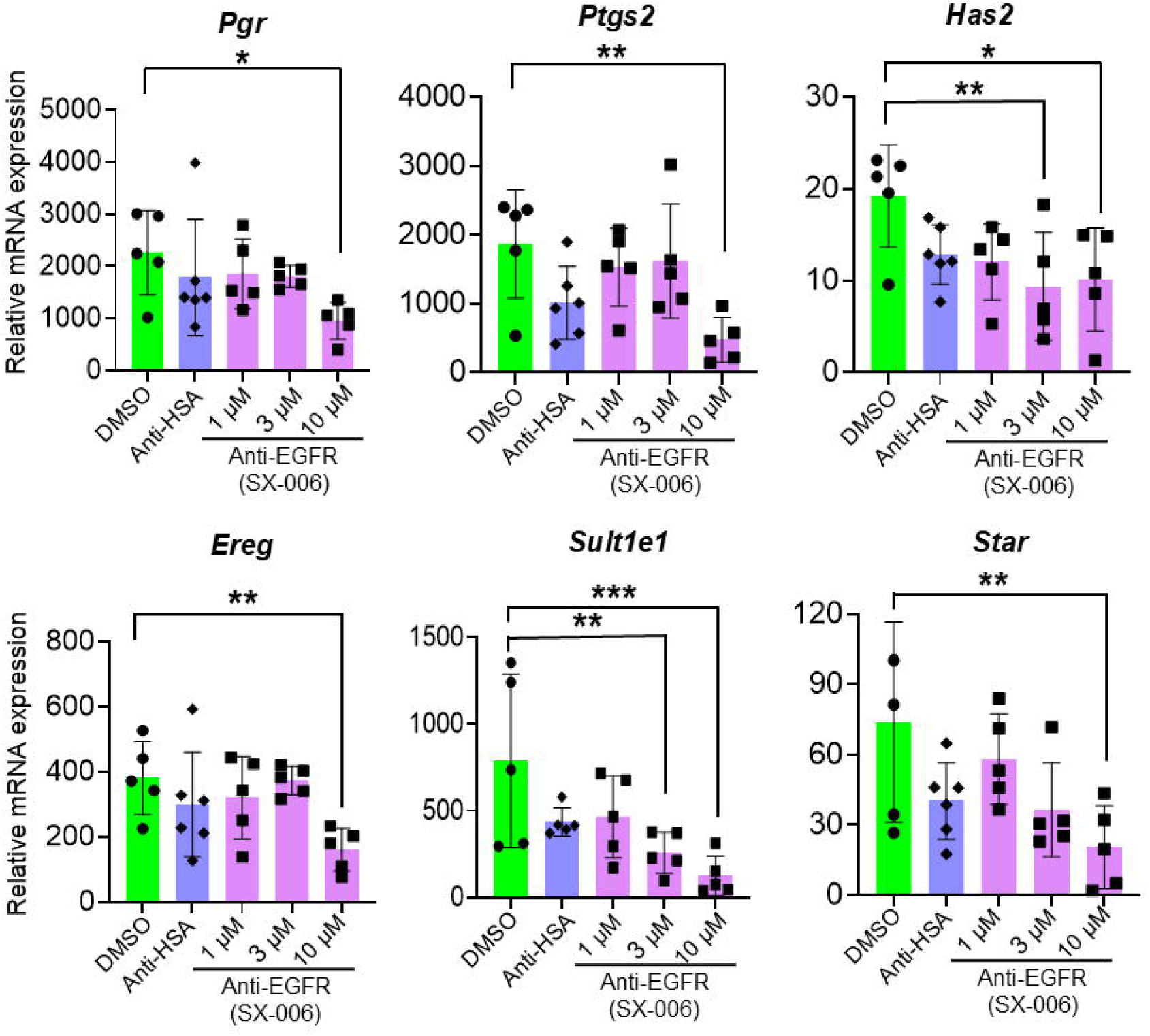
SX-006 attenuates hCG-induced ovulatory gene expression. Follicles were treated with DMSO, anti-HSA DARPin, or anti-EGFR DARPin SX-006 at 1, 3, or 10 μM during hCG-induced ovulation and collected at 4 h post-hCG. Ovulation-related gene expression was measured by single-follicle RT-qPCR. Gene expression levels were normalized to the corresponding pre-hCG follicles, which were set to 1. Data are presented as mean ± SEM. n = 5–6 follicles per group. Statistical significance was determined using one-way ANOVA followed by Dunnett’s multiple comparisons test. Asterisks indicate significant differences compared with the DMSO control group. \**P* < 0.05, \*\**P* < 0.01, \*\*\**P* < 0.001.

### Anti-EGFR DARPin SX-006 disrupts ovulation-related transcriptional programs

To further elucidate the molecular mechanisms of Anti-EGFR DARPin-induced anovulation in an unbiased manner, follicles treated with vehicle and 10 μM SX-006 at 4 h post-hCG treatment were collected for single-follicle RNA-seq analysis. Principal component analysis (PCA) revealed clear separation between control and SX-006-treated follicles, indicating distinct transcriptional profiles in response to SX-006 treatment (Figure 5A). There were 1219 differentially expressed genes (DEGs) with fold change >= 2 or <= 0.5 and FDR < 0.05, including 706 up- and 513 down-regulated genes induced by SX-006 (Figure 5B). The top 10 genes in each direction are highlighted in the volcano plot in Figure 5B, and all DEGs were listed in Table S1.

**Figure 5.**
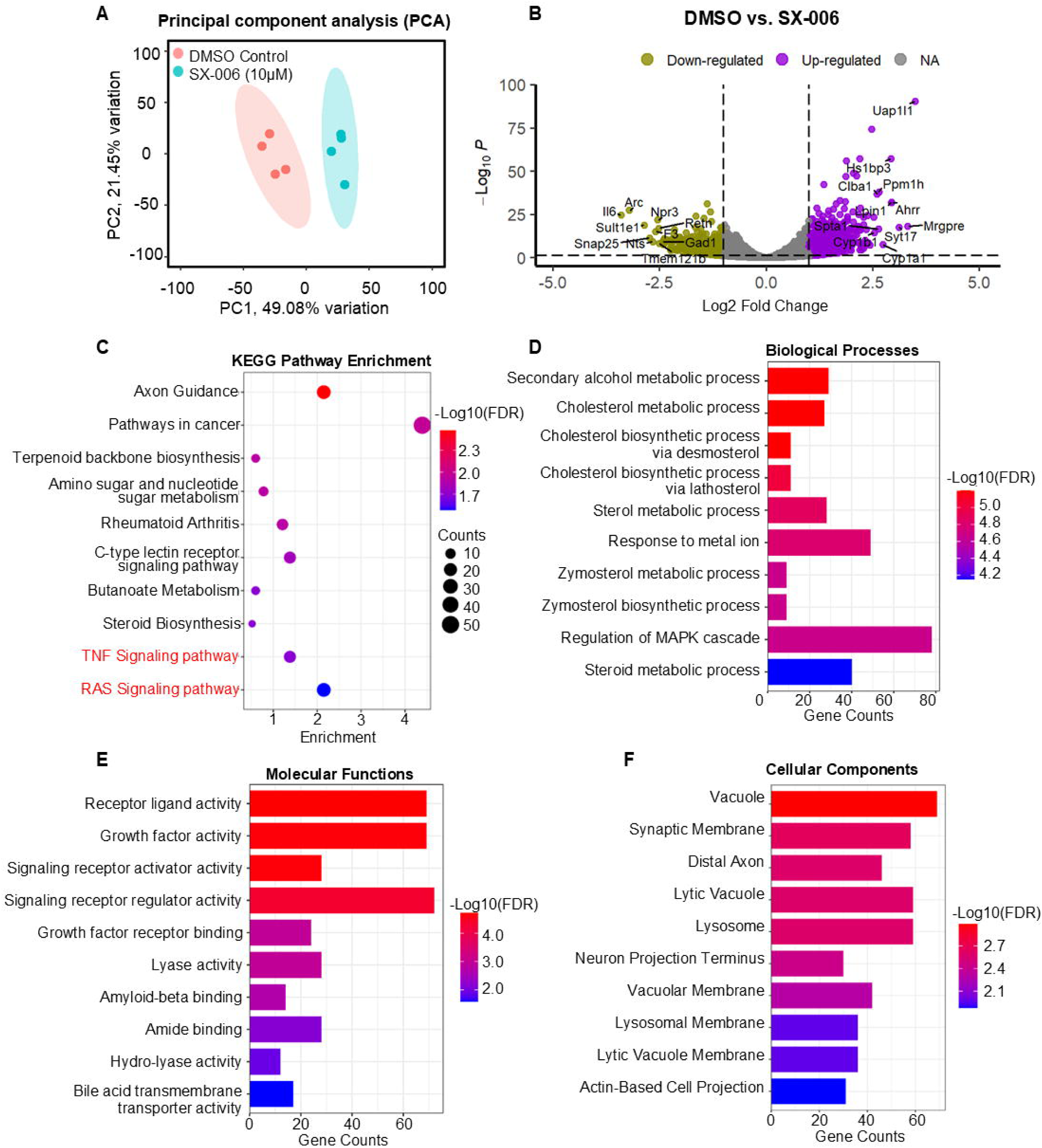
**SX-006 broadly disrupts the hCG-induced ovulatory transcriptome**. Follicles were treated with DMSO or 10 μM SX-006 during hCG-induced ovulation and collected at 4 h post-hCG for single-follicle RNA-seq (n = 4). (A) Principal component analysis (PCA) of transcriptomic profiles showing separation between control and SX-006-treated follicles. (B) Volcano plot showing differentially expressed genes (DEGs) identified between control and SX-006-treated follicles. Upregulated and downregulated genes are highlighted in purple and olive, respectively. (C–F) KEGG pathway and GO enrichment analyses of DEGs, including the top 10 enriched KEGG pathways (C), biological processes (D), molecular functions (E), and cellular components (F). Bubble size in the KEGG plot indicates gene count, and color represents enrichment significance. The x-axis in the GO enrichment plots represents the number of DEGs associated with each pathway or GO term. Bar colors indicate enrichment significance expressed as −log10(FDR).

DEGs were subjected to GO and KEGG enrichment analyses. All DEGs and significantly enriched GO terms and pathways are provided in Excel Table S2-S3, and the top 10 enriched terms in each category are shown in Figure 5C–5F. GO biological process analysis identified significant enrichment of genes involved in “Secondary Alcohol Metabolic Process”, “Cholesterol Metabolic Process”, and “Cholesterol Biosynthetic Process via Desmosterol”. Molecular function analysis revealed enrichment of “Receptor Ligand Activity”, “Growth Factor Activity”, and “Signaling Receptor Activator Activity”. Cellular component analysis highlighted enrichment of “Vacuole”, “Synaptic Membrane”, and “Distal Axon”. KEGG pathway analysis demonstrated significant enrichment of “Axon Guidance”, “Pathways in cancer”, and “Terpenoid Backbone Biosynthesis” pathways.

To further characterize these transcriptomic changes, GO enrichment analysis was conducted using SX-006-induced downregulated genes. All significantly enriched GO terms and pathways are listed in Excel Table S4, and the top 10 enriched terms in each category are shown in Figure S3. GO biological process analysis identified significant enrichment of genes associated with “Ovulation Cycle”, “Vascular Process in Circulatory System”, and “Regulation of MAPK Cascade”. Molecular function analysis revealed enrichment of “Receptor Ligand Activity”, “Signaling Receptor Activator Activity” and “Signaling Receptor Regulator Activity”. Cellular component analysis highlighted enrichment of “Neuron Projection Terminus”, “Receptor Complex”, and “Postsynaptic Membrane”. To further examine the suppression of core ovulation-associated genes by SX-006, the expression of major ovulatory marker genes were visualized by heatmap analysis (Figure S3). Consistent with the RT-qPCR results, SX-006 broadly attenuated the expression of genes involved in EGFR signaling, cumulus expansion, follicle rupture, and steroidogenesis.

### Distinct EGFR perturbation modes induce distinct transcriptomic changes

To further elucidate the mechanisms of extracellular EGFR blockade and intracellular EGFR kinase inhibition in ovulation perturbation, we compared DEGs identified in follicles treated with anti-EGFR DARPin (SX-006) or with small molecule AG1478-treated follicles at 4 h post-hCG treatment. The RNA-seq data from AG1478-treated follicles were generated in an independent experimental batch in our laboratory, using the same experimental and sequencing procedures. SX-006 treatment produced a larger transcriptional response than AG1478, with 1,096 SX-006-specific DEGs, 453 AG1478-specific DEGs, and 123 overlapped DEGs (Figure 6A). To further characterize these transcriptomic differences, GO enrichment analysis was performed separately on SX-006-specific, AG1478-specific, and shared DEGs. All significantly enriched GO terms are listed in Excel Tables S5–S7. Functional enrichment analysis revealed distinct biological processes associated with each DEG set. The shared DEGs were enriched for processes related to “Sterol Biosynthetic Process” “Ovulation Cycle”, “Zymosterol Biosynthetic Process”, “Vasculature development” and “ Cholesterol Biosynthetic

**Figure 6.**
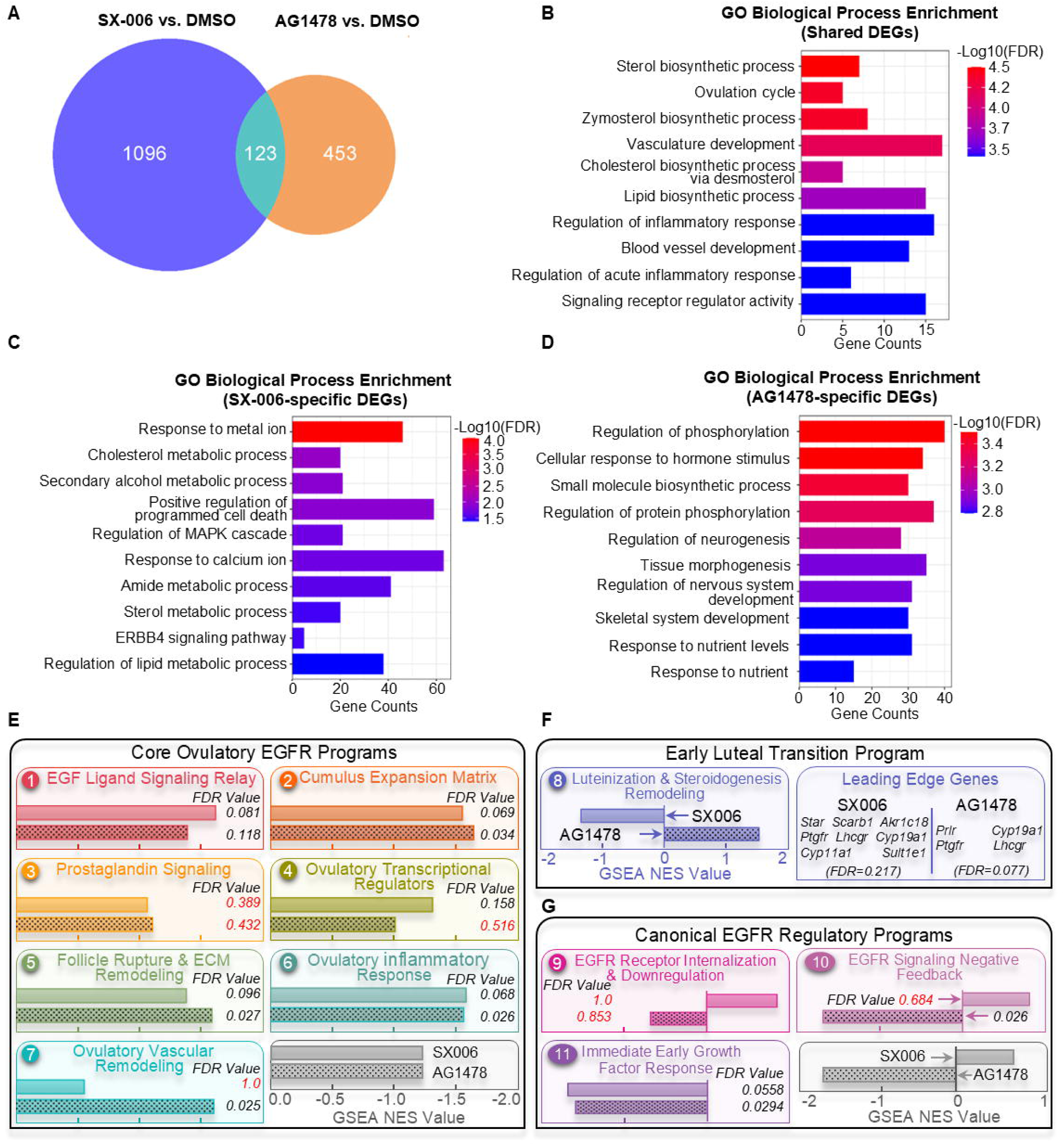
SX-006 and Tyrphostin AG1478 induce overlapping but distinct transcriptional programs in periovulatory follicles. (A) Venn diagram of DEGs from SX-006-treated follicles(n = 4) and Tyrphostin AG1478(n=3) dataset collected 4 h post-hCG. (B–D) GO Biological Process enrichment analysis of shared DEGs (B), SX-006-specific DEGs (C), and AG1478-specific DEGs (D). Bar length indicates gene count and color indicates enrichment significance [−log10(FDR)]. (E) Normalized enrichment scores (NES) and FDR q values of core ovulatory EGFR programs, including EGF ligand signaling relay, cumulus expansion matrix, prostaglandin signaling, ovulatory transcriptional regulators, follicle rupture and extracellular matrix (ECM) remodeling, ovulatory inflammatory response, and ovulatory vascular remodeling. (F) NES and leading-edge genes of the luteinization and steroidogenic remodeling module of early luteal transition program. (G) NES and FDR q values of canonical EGFR regulatory programs, including EGFR receptor internalization and downregulation, EGFR signaling negative feedback, and immediate-early growth factor response modules. Positive NES values indicate relative enrichment, and negative NES values indicate relative suppression of the corresponding gene set. FDR q values < 0.25 were considered significant enrichment according to GSEA criteria; modules with FDR q values ≥ 0.25 are indicated in red. Follicles were collected 4 h after hCG stimulation for RNA-seq analysis.

Process via Desmosterol” (Figure 6B). DEGs specific to SX-006 were primarily related to “Response to Metal Ion”, “Cholesterol Metabolic Process”, “Secondary Alcohol Metabolic Process”, “Positive Regulation of Programmed Cell Death” and “Regulation of MAPK Cascade”, whereas AG1478-specific DEGs were associated with “Regulation of Phosphorylation”, “Cellular Response to Hormone Stimulus”, “Small Molecule Biosynthetic Process”, “Regulation of Protein Phosphorylation”, and “Regulation of Neurogenesis” (Figure 6B). These results suggest that SX-006 and AG1478 perturb overlapping but also distinct follicular transcriptional programs downstream of the EGFR signaling.

### Extracellular EGFR blockade and intracellular kinase inhibition converge on ovulatory programs but diverge in EGFR regulatory and steroidogenic remodeling responses

To further compare extracellular EGFR blockade and intracellular kinase inhibition at the pathway level, we performed GSEA using 12 literature-curated EGFR-associated modules representing core ovulatory EGFR programs, oocyte-associated program, early luteal transition programs, and canonical EGFR regulatory programs (Figure S4-S6, Excel Table S7-S11) [19, 24–36]. Overall, SX-006 and AG1478 exhibited broadly similar enrichment patterns across the core ovulatory EGFR programs (Figure 6E). Both treatments negatively enriched the EGF ligand signaling relay, cumulus expansion matrix, prostaglandin signaling, ovulatory transcriptional regulators, follicle rupture and extracellular matrix remodeling, ovulatory inflammatory response, and ovulatory vascular remodeling modules. Significant negative enrichment (FDR < 0.25) was observed in both treatment groups for the EGF ligand signaling relay, cumulus expansion matrix, follicle rupture and extracellular matrix remodeling, ovulatory inflammatory response, and immediate-early growth factor response modules. The ovulatory transcriptional regulators module reached significant enrichment only in SX-006-treated follicles, whereas the ovulatory vascular remodeling module reached significant enrichment only in AG1478-treated follicles. In contrast, the prostaglandin signaling module did not reach the predefined significance threshold in either comparison. Together, these findings indicate that extracellular EGFR blockade and intracellular kinase inhibition similarly suppress a common EGFR-dependent ovulatory transcriptional program.

A notable difference between SX-006 and AG1478 emerged in the luteinization and steroidogenic remodeling module (Figure 6F). SX-006 negatively enriched this module (NES = −1.35, FDR = 0.22), whereas AG1478 positively enriched the same module (NES = 1.55, FDR = 0.08). Further leading-edge gene analysis revealed distinct contributing genes between treatments. In SX-006-treated follicles, negative enrichment was primarily driven by *Star, Scarb1, Akr1c18, Cyp11a1*, and *Sult1e1*, genes associated with cholesterol utilization, steroidogenic capacity, and progesterone biosynthesis. In contrast, AG1478-positive enrichment was driven by *Prlr, Lhcgr, Ptgfr,* and *Cyp19a1*, genes more closely associated with endocrine responsiveness and the granulosa cell state. These findings suggest that extracellular EGFR blockade and intracellular kinase inhibition differentially influence early follicle-to-luteal transition programs. Specifically, SX-006 was associated with suppression of the steroidogenic machinery during the early stages of luteinization, consistent with the reduced progesterone secretion observed at later time points (Figure 3F). In contrast, the positive enrichment observed in AG1478-treated follicles was not driven by core steroidogenic machinery genes, suggesting that the effects of AG1478 on progesterone production in Figure 3F may occur through later-stage, indirect, or non-transcriptional regulatory mechanisms.

Differences were also observed in canonical EGFR regulatory programs (Figure 6G). AG1478 significantly negatively enriched the EGFR signaling negative feedback module (NES = −1.71, FDR = 0.026), whereas SX-006 did not. Leading-edge genes contributing to AG1478 enrichment included *Dusp4*, *Dusp6*, *Errfi1*, and *Spry2*, which are established components of the EGFR feedback network. These findings suggest that intracellular kinase inhibition suppresses transcriptional programs that attenuate EGFR signaling more strongly. In contrast, the EGFR receptor internalization and downregulation module was not significantly enriched in either treatment group, although opposite enrichment directions were observed between SX-006 and AG1478. Because this module contains genes involved in receptor internalization and endosomal sorting, including *Hgs*, *Stam*, *Eps15*, and *Epn1*, these results raise the possibility that extracellular EGFR blockade and intracellular kinase inhibition may differentially influence receptor handling and receptor-associated regulatory processes.

### Anti-EGFR DARPin SX-006 induces AhR target gene expression

RNA-seq analysis unexpectedly revealed that anti-EGFR DARPin SX-006 induced the expression of several aryl hydrocarbon receptor (AhR)-responsive genes during hCG-induced ovulation, including *Ahrr*, *Cyp1a1*, and *Cyp1b1*, which were among the top 10 most upregulated genes in SX-006-treated follicles (Figure 5B). AhR is a ligand-activated transcription factor that can be activated by diverse endogenous ligands, such as kynurenine and tryptophan metabolites, and xenobiotics including dioxins and polycyclic aromatic hydrocarbons (PAHs) [52–54]. Upon ligand binding, AhR heterodimerizes with the AhR nuclear translocator (ARNT) and binds to xenobiotic response elements (XREs), leading to the transcriptional activation of canonical target genes, including *Cyp1a1* [55–57]. CYP1A1-mediated metabolism has been shown to further generate reactive intermediates and reactive oxygen species (ROS), thereby contributing to oxidative stress responses [58–62].

To determine whether the induction of AhR-responsive genes was specific to extracellular EGFR targeting or caused by the general DARPin scaffold, follicles treated with anti-HSA DARPin, AG1478, or anti-EGFR DARPin SX-006 were collected at 4 h post-hCG treatment for RT-qPCR analysis. The results showed that *Ahrr* and *Cyp1a1* expression levels remained comparable among vehicle-, AG1478, and anti-HSA DARPin-treated follicles, whereas SX-006 significantly induced their expression (Figure 7). In contrast, oxidative stress-related genes, including *Sod1* and *Sod2*, exhibited no significant changes among all treatment groups (Figure 7). Collectively, these findings suggest that extracellular EGFR blockade by anti-EGFR DARPin SX-006 uniquely induces AhR-responsive transcriptional programs during ovulation, although the underlying mechanism remains elusive.

**Figure 7.**
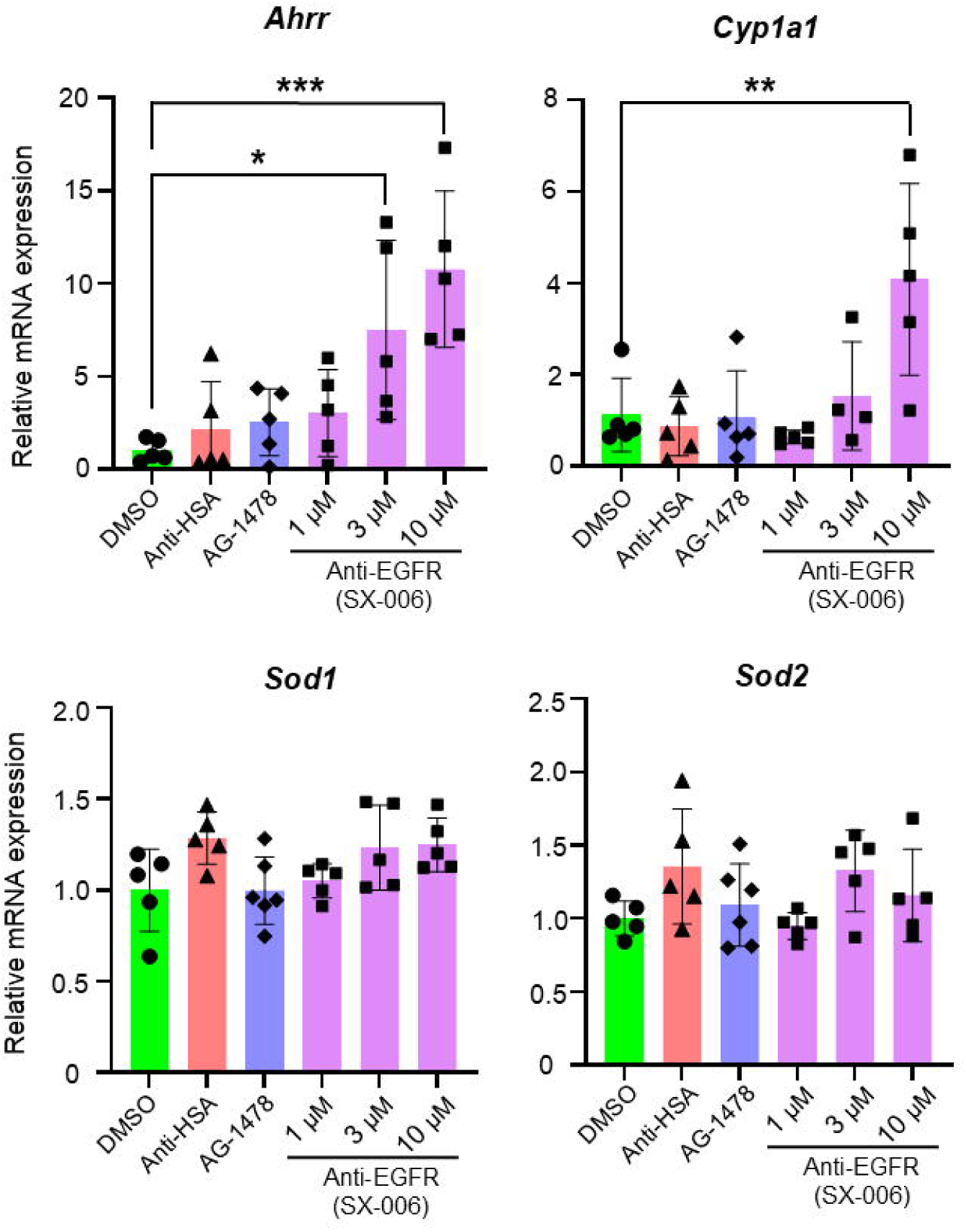
Anti-EGFR DARPin SX-006 induces AhR-responsive gene expression during hCG-induced ovulation. Follicles were treated with vehicle control, anti-HSA DARPin, Tyrphostin AG-1478, or anti-EGFR DARPin SX-006 during hCG-induced ovulation and collected at 4 h post-hCG. Expression levels of the AhR-responsive genes *Ahrr* and *Cyp1a1*, as well as the oxidative stress-related genes *Sod1* and *Sod2*, were measured by RT-qPCR. Data are presented as mean ± SEM. n = 5–6 follicles per group. Statistical significance was determined using one-way ANOVA followed by Dunnett’s multiple comparisons test. Asterisks indicate significant differences compared with the DMSO control group. \**P* < 0.05, \*\**P* < 0.01, \*\*\**P* < 0.001.

## DISCUSSION

Mechanistic studies of ovarian biology remain challenging due to the lack of highly selective, tunable, and physiologically relevant tools. Existing approaches, including genetic animal models, small molecules, and antibodies, each possess limitations [32, 63, 64]. Using EGFR as a proof-of-concept target in combination with our IVFG-based *ex vivo* ovulation system, our study establishes DARPin as an efficient, versatile, and tunable platform for studying ovarian signaling. We demonstrate that anti-EGFR DARPins, particularly the bispecific SX-006 that incorporates a central leucine zipper motif adapted from E69-LZ3-E01 to drive homodimerization, selectively inhibit follicle ovulation without overt cytotoxicity. Moreover, extracellular EGFR blockade produced transcriptional and functional effects distinct from intracellular EGFR kinase small molecule inhibitor, suggesting that extracellular ligand–receptor perturbation and intracellular kinase inhibition represent biologically distinct modes of EGFR modulation during ovulation. Together, these findings support the utility of DARPins not only as experimental tools for dissecting ovarian biology but also as a potential therapeutic modality for ovarian disorders, infertility, and non-hormonal contraception.

EGFR has been well established as a central mediator of LH/hCG-induced ovulation. Prior studies using genetically modified mouse models and small-molecule EGFR inhibitors demonstrated that EGFR activation is required for cumulus expansion, oocyte maturation, and follicle rupture [19–21]. However, these studies primarily disrupt receptor expression or intracellular kinase activity and do not specifically interrogate extracellular ligand–receptor interactions. Here, we used anti-EGFR DARPins to selectively target the extracellular ligand-binding interface of EGFR. SX-006 potently inhibited follicle rupture and suppressed ovulation-associated gene expression, supporting the concept that extracellular EGFR activation is required for proper ovulatory signaling. Importantly, these findings demonstrate the feasibility of using engineered DARPins to modulate ovarian signaling pathways within intact follicles.

One important finding of this study is that extracellular EGFR blockade and intracellular EGFR kinase inhibition produced both overlapping and distinct transcriptional responses in periovulatory follicles. Consistent with the central role of EGFR signaling in ovulation, SX-006 and AG1478 converged on a common EGFR-dependent ovulatory transcriptional program, suppressing modules associated with EGF ligand signaling, cumulus expansion, follicle rupture, inflammatory responses, and immediate-early transcriptional activation. However, notable differences emerged in pathways related to early luteal transition and EGFR regulatory responses. SX-006 preferentially suppressed genes associated with cholesterol utilization and steroidogenic remodeling, whereas AG1478 positively enriched a distinct set of genes, including *Prlr*, *Lhcgr*, *Ptgfr*, and *Cyp19a1*. In addition, AG1478 suppressed canonical EGFR negative feedback regulators more strongly, including *Dusp4*, *Dusp6*, *Errfi1*, and *Spry2*. Together, these findings suggest that although extracellular receptor blockade and intracellular kinase inhibition similarly disrupt ovulatory signaling, they differentially influence steroidogenic remodeling and EGFR regulatory networks, highlighting partially distinct downstream transcriptional consequences of targeting EGFR at different levels of the signaling cascade.

Our *ex vivo* ovulation testing results reveal that follicle rupture is more sensitive to extracellular EGFR blockade than oocyte meiotic maturation or progesterone secretion. Lower concentrations of SX-006 inhibited follicle rupture without substantially affecting the first polar body extrusion or luteinization, whereas higher concentrations disrupted multiple ovulatory endpoints. In comparison, the small-molecule EGFR inhibitor AG1478 produced broader suppression of all ovulatory functions. These findings suggest that distinct ovulatory processes may require different thresholds, durations, or modes of EGFR signaling. Follicle rupture is a highly coordinated inflammatory-like process involving prostaglandin synthesis, cytokines, ECM remodeling, and weakening of the follicle wall. Partial disruption of extracellular ligand-dependent EGFR amplification by anti-EGFR DARPin may preferentially impair follicle rupture before broader effects on meiotic progression and luteinization become evident. In contrast, intracellular kinase inhibition by AG1478 is likely to produce more widespread suppression of downstream signaling cascades, thereby affecting multiple ovulatory endpoints simultaneously. Together, these observations support the concept that extracellular receptor blockade and intracellular kinase inhibition represent biologically distinct modes of EGFR perturbation during ovulation.

An unexpected finding from this study was the anti-EGFR DARPin-induced AhR-responsive genes, including *Ahrr*, *Cyp1a1*, and *Cyp1b1*. Our results further demonstrate that this response was not observed in follicles treated with anti-HSA DARPin or AG1478, suggesting that the induction was not attributable to the general DARPin scaffold. Although the underlying mechanism remains unclear, emerging evidence suggests extensive crosstalk between EGFR and AhR signaling pathways in multiple tissues [65, 66]. AhR interacts with growth factor signaling networks through shared downstream mediators, transcriptional co-regulators, inflammatory signaling pathways, and receptor trafficking processes [67–69]. As SX-006 specifically targets the extracellular EGFR ligand-binding interface, one possibility is that extracellular receptor blockade alters membrane-associated signaling dynamics or compensatory ligand-responsive pathways that secondarily activate AhR-responsive transcriptional programs. Intriguingly, AhR activation has also been implicated in ovulatory dysfunction and ovarian toxicology, including dioxin- and PAH-induced inhibition of ovulation [70]. Therefore, the induction of AhR-responsive genes may reflect a previously unrecognized adaptive or stress-responsive signaling associated with extracellular EGFR perturbation during ovulation.

A broader significance of this study extends beyond EGFR biology. DARPin possesses several features that make it a particularly attractive tool for studying reproductive biology, including small molecular size, high stability, modular engineering flexibility, tunable valency, and extracellular receptor specificity. Compared with traditional monoclonal antibodies, DARPins lack Fc-mediated immune effector functions and can be efficiently produced in bacterial systems [8]. Compared with small-molecule inhibitors, DARPins provide a fundamentally different mode of pathway perturbation by targeting extracellular receptors rather than intracellular catalytic inhibition [71]. This extracellular targeting strategy may be especially useful for studying ligand-dependent autocrine and paracrine signaling pathways that coordinate communication among granulosa cells, cumulus cells, and oocytes.

Several limitations should be considered. For example, the anti-EGFR DARPins we used were originally engineered against human EGFR, whereas the functional assays were conducted using murine follicles. Although SX-006 exhibited the strongest murine EGFR binding among the tested constructs and produced clear biological effects, its affinity toward murine EGFR remained weaker than toward human EGFR. Future development of murine-specific or humanized EGFR-targeting DARPins will strengthen this platform. Moreover, although transcriptomic and functional analyses strongly support altered EGFR-associated signaling, direct measurements of receptor phosphorylation, receptor internalization, and downstream ERK activation will be crucial to define the precise signaling mechanisms induced by extracellular EGFR blockade.

In summary, this study establishes DARPins as a novel molecular tool for studying ovarian signaling. Using EGFR as a proof-of-concept target, we demonstrate that extracellular EGFR blockade inhibits ovulation and induces transcriptional programs distinct from those induced by intracellular kinase inhibition. These findings support the concept that extracellular receptor perturbation represents a biologically distinct mode of pathway modulation within the ovary. More broadly, this work introduces DARPins as a versatile, tunable platform for investigating ovarian biology and reproductive signaling networks, and highlights their potential for studying ovarian disorders, infertility, and the development of next-generation non-hormonal contraceptives.

## AUTHOR CONTRIBUTIONS

Y. Liu contributed to experimental design, follicle isolation, *ex vivo* follicle culture, hCG-induced ovulation assays, data collection, data analysis, result interpretation, qPCR validation, statistical analysis, and manuscript drafting. Y. Liu, C. Mitra, and Yue Liu contributed to the preparation, purification, validation, and related data interpretation of the DARPin protein. J. Zhang, Y. Liu, S. Liu, H. M. VanBenschoten, and B. Goods conducted RNA-seq data processing, differential expression analysis, pathway enrichment analysis, and data visualization, and contributed to AG1478 treatment experiments, sequencing sample preparation, and related data collection. F. Chen contributed to experimental design, DARPin vector design, manuscript editing, and critical revision. F. Chen and S. Xiao conceived and supervised the study, designed experiments, analyzed and interpreted data, wrote and revised the manuscript, and approved the final manuscript.

## FUNDING

This work was supported by the National Institutes of Health (NIH) R01ES032144 and R01HD115810 to S. Xiao. Tyrphostin AG1478-related RNA-seq data was supported in part by the Gates Foundation (INV-003385) to S. Xiao and B. A. Goods. Under the grant conditions of the Gates Foundation, a Creative Commons Attribution 4.0 Generic License has already been assigned to the Author Accepted Manuscript version that might arise from this submission.

## Grant support

This work was supported by the National Institutes of Health (NIH) R01ES032144 and R01HD115810 to S. Xiao. Tyrphostin AG1478-related RNA-seq data was supported in part by the Gates Foundation (INV-003385) to S. Xiao and B. A. Goods. Under the grant conditions of the Gates Foundation, a Creative Commons Attribution 4.0 Generic License has already been assigned to the Author Accepted Manuscript version that might arise from this submission.

## Declaration of interest

The authors declare no conflict of interest.

---

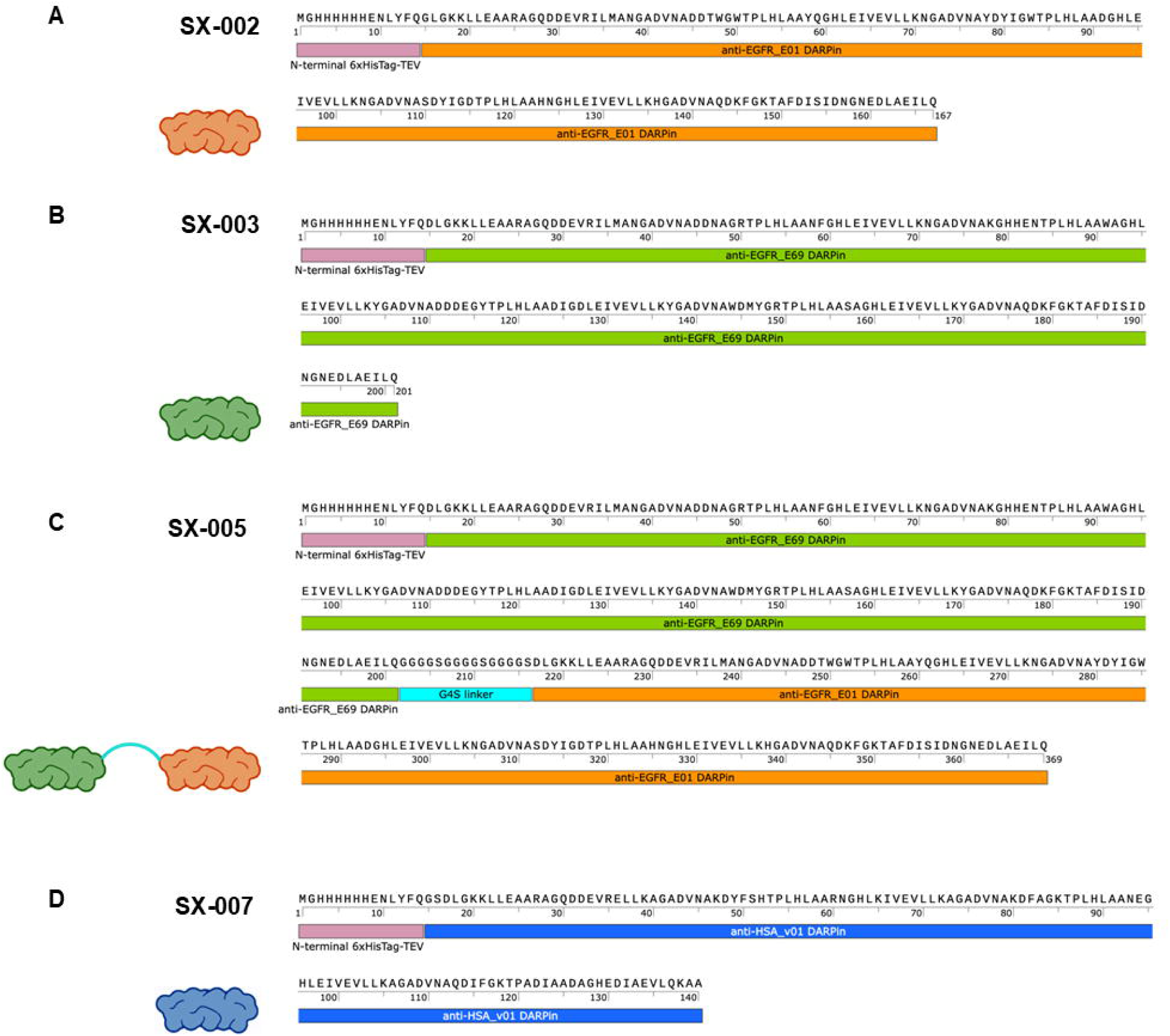

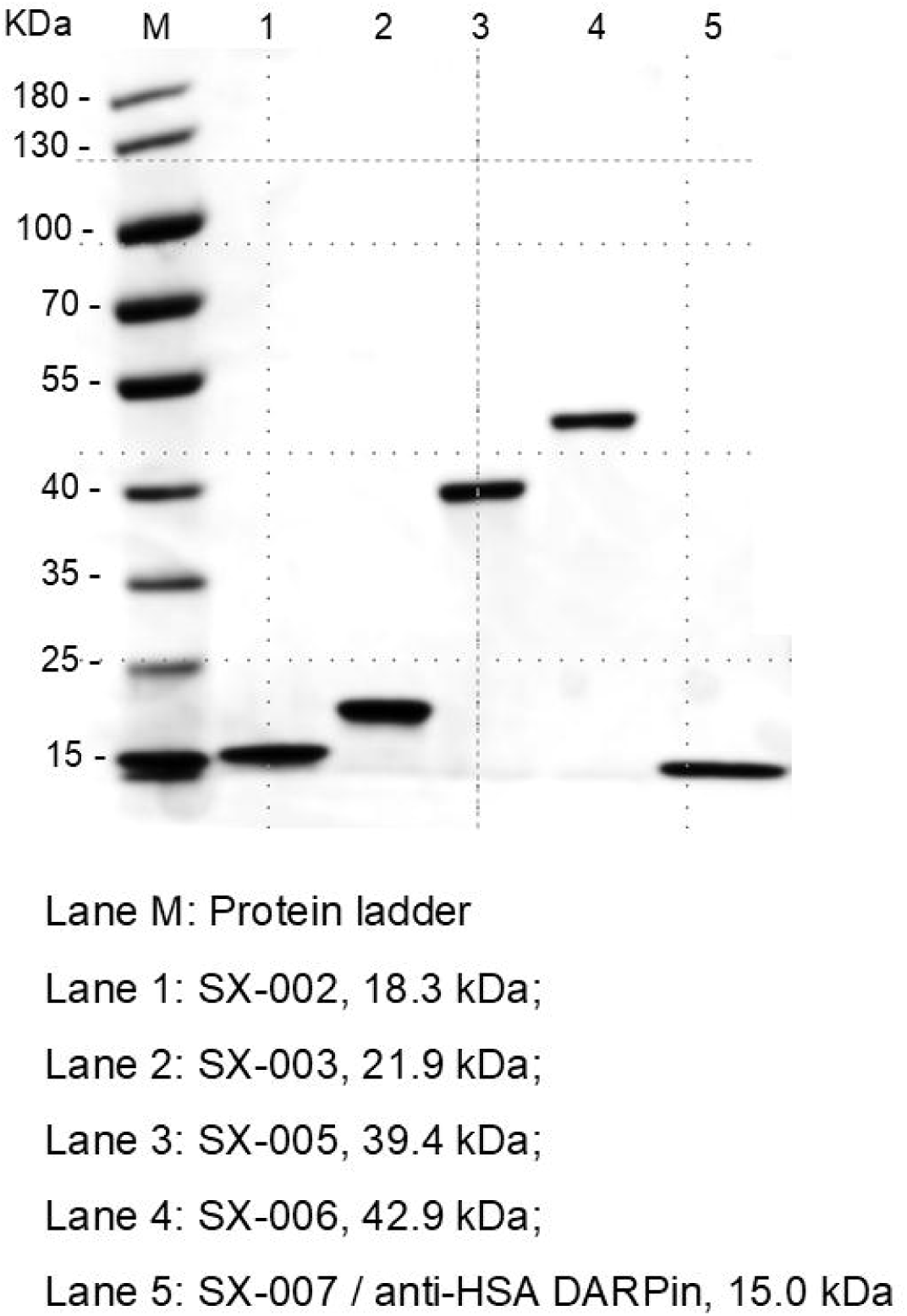

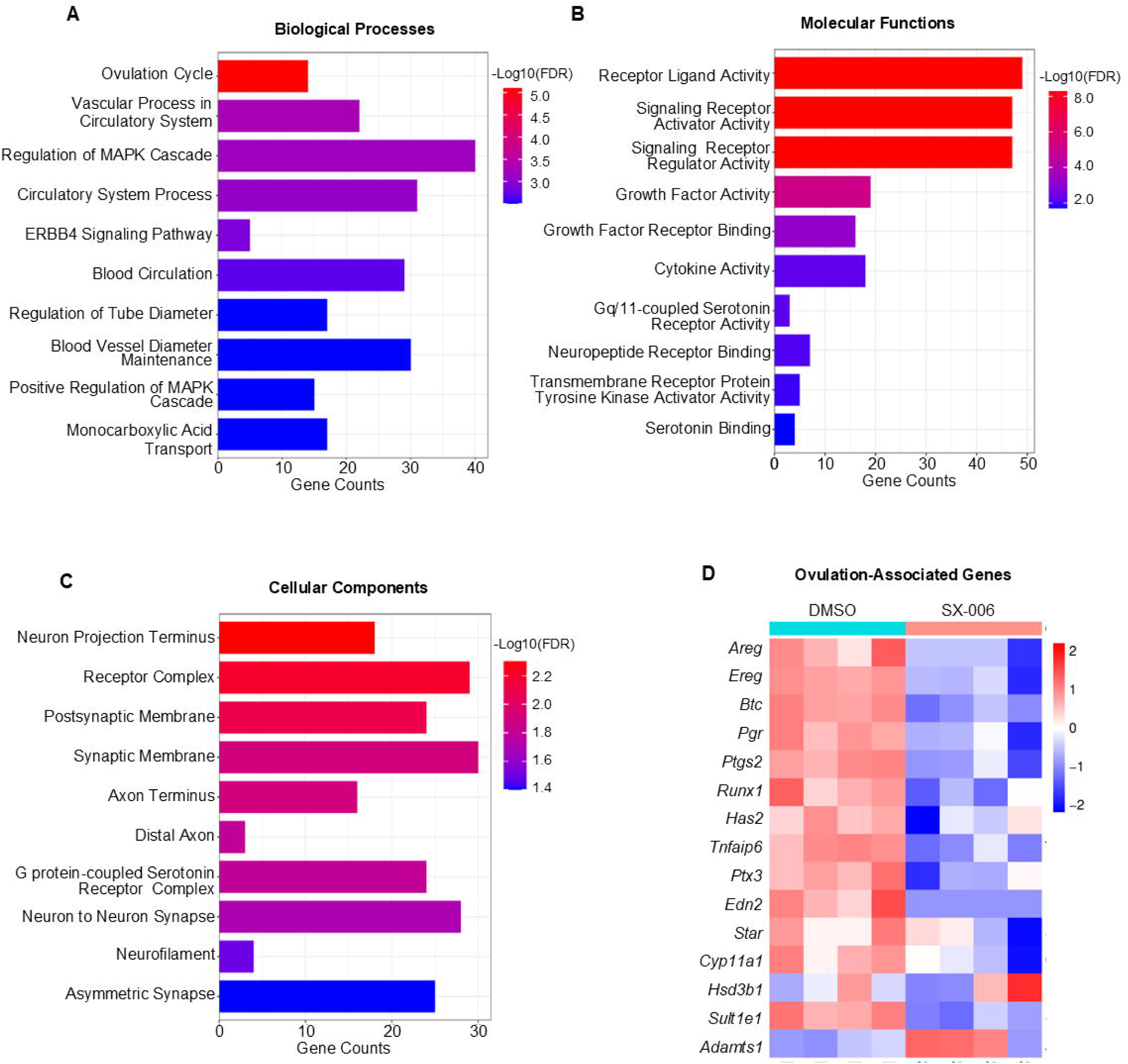

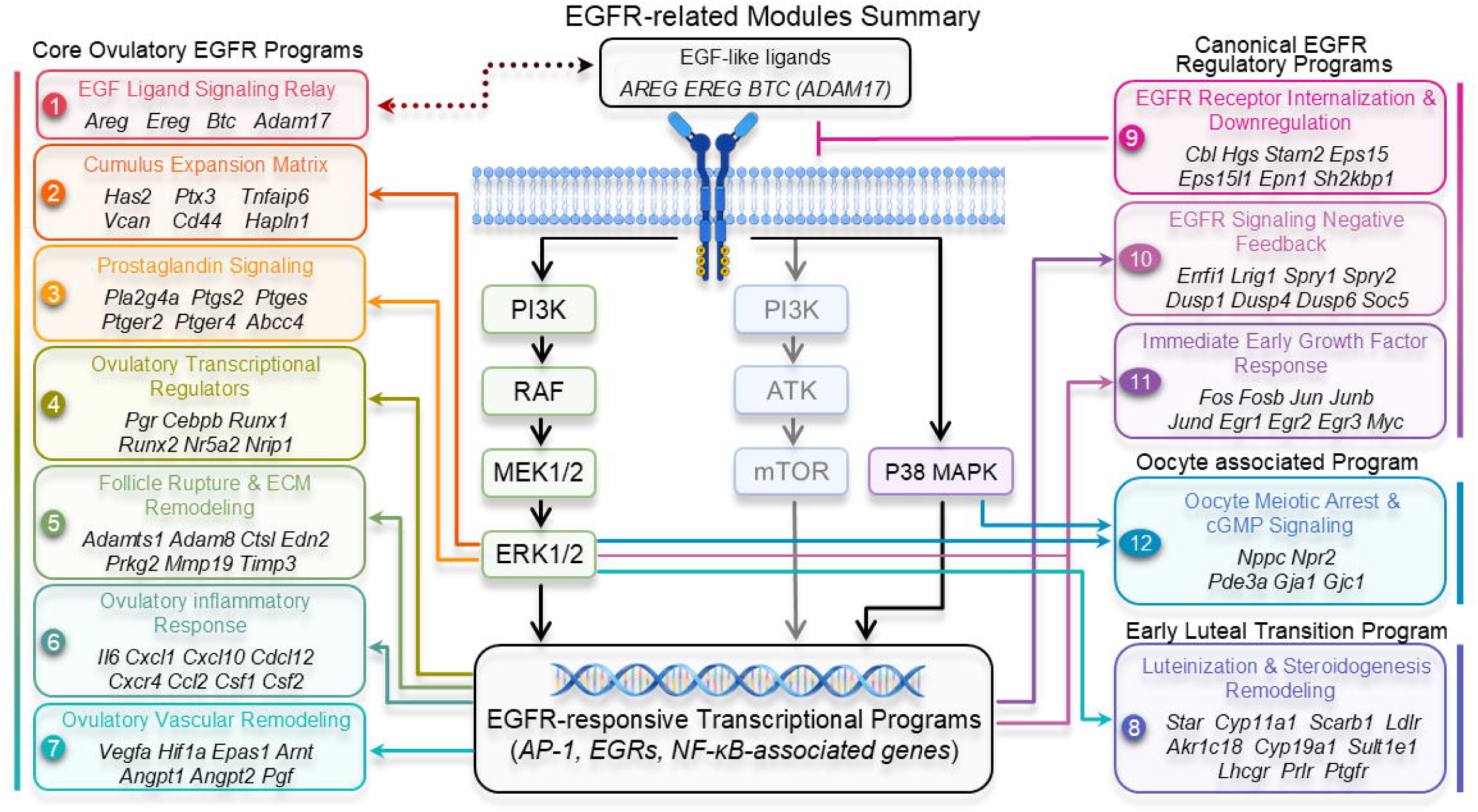

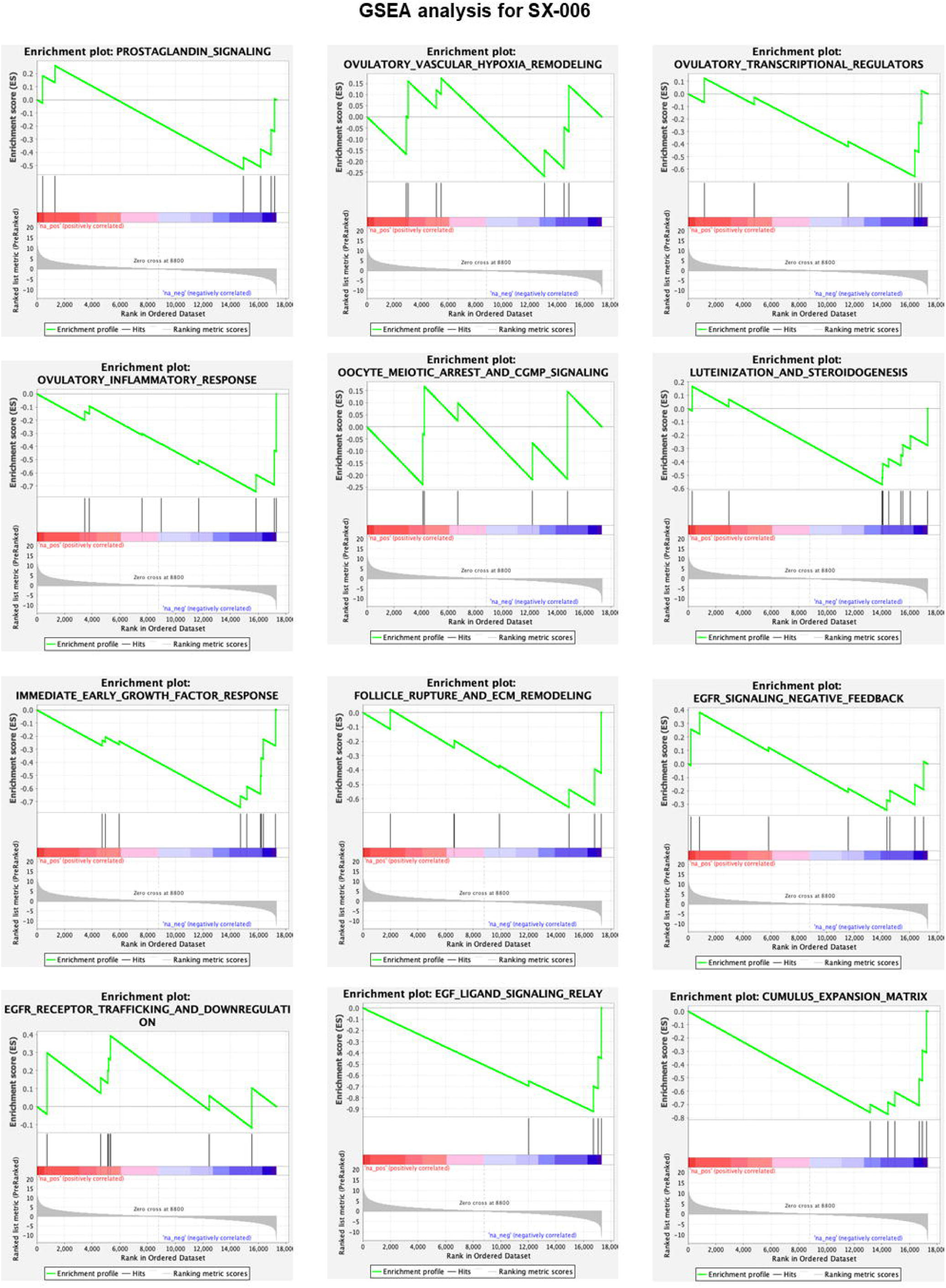

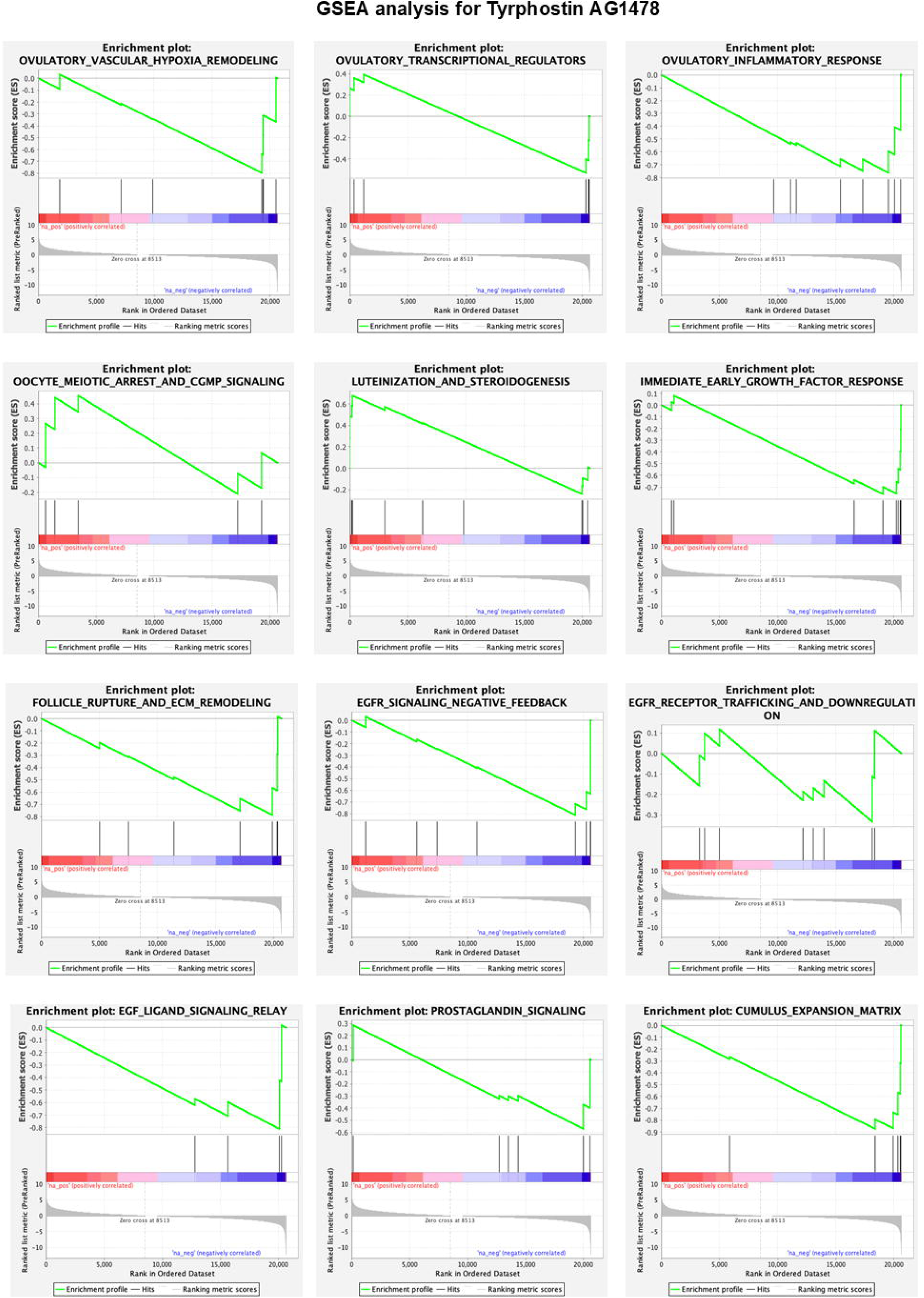

## Notes

### Competing Interest Statement

The authors have declared no competing interest.

