## Supplementary material for "A Designed Ankyrin Repeat Protein (DARPin) Targeting EGFR Inhibits Ovulation and Enables a Novel Platform for Studying Ovarian Biology and Pathophysiology": Figure S

**Table S1.** Expression scale and protein yield of anti-EGFR DARPins. Summary of recombinant DARPin protein yields (mg/L) obtained from *E. coli* expression batches (30 mL-scale).

| Name | MW | Extinction coefficient (ε) @280 nm | Charge at pH 7 | pI (isoelectric point) | l* | Protein yields |
| --- | --- | --- | --- | --- | --- | --- |
|  | kDa | (mg/mL)^-1^ cm^-1^ |  |  | cm | mg/L |
| SX-002 | 18.3 | 1.283 | -14.7 | 5.10 | 1.0 | 50.71 |
| SX-003 | 21.9 | 0.911 | -18.4 | 4.97 | 1.0 | 69.26 |
| SX-005 | 39.4 | 1.077 | -37.7 | 4.62 | 1.0 | 108.90 |
| SX-006 | 42.9 | 0.991 | -36.6 | 4.76 | 1.0 | 80.73 |
| SX-007 | 15.0 | 0.171 | -6.8 | 5.78 | 1.0 | 69.60 |

**Table S2.** Primer sequences of genes by RT-qPCR, including ovulatory marker genes, AhR target genes, and oxidative stress related genes.

| **Gene name** | **Sequence** |
| --- | --- |
| *Pgr* | F: ATGGTCCTTGGAGGTCGTAA  R: GTGGCGGGACCAGTTGAAT |
| *Ptgs2* | F: TGGGGGAAGAAATGTGCCAA  R: CAGCCATTTCCTTCTCTCCTGT |
| *Has* | F: GCCGGTCGTCTCAAATTCATC  R: ACCTCTCACAATGCATCTTGTTC |
| *Ereg* | F: GCATCCCAGGAGAATCCGAG  R: GTGTAGCCCACTTCACATCTGC |
| *Sult1e1* | F: CTTCCAGGAGATGAAGAACAATCC  R: GGAAGTGGTTCTTCCAGTCTCC |
| *Star* | F: TTGGGCATACTCAACAACCA  R: CCTTGACATTTGGGTTCCAC |
| *Hprt* | F: TCAGTCAACGGGGGACATAAA  R: GGGGCTGTACTGCTTAACCAG |
| *Ahrr* | F: GTTGGATCCTGTAGGGAGCA  R: AGTCCAGAGGCTCACGCTTA |
| *Cyp1a1* | F: TCTCGTGGAGCCTCATGTACCT  R: TGCCGATCTCTGCCAATCA |
| *Cyp1b1* | F: TTGACCCCATAGGAAACTGC  R: GCTGTCTCTTGGTAGGAGGA- |
| *Sod1* | F: GGTGAACCAGTTGTGTTGTCAGG  R: ATGAGGTCCTGCACTGGTACAG |
| *Sod2* | F: TAACGCGCAGATCATGCAGCTG  R: AGGCTGAAGAGCGACCTGAGTT |

*F: forward; R: reverse

**Table S3.** Genes examined by RT-qPCR and their reported functions in ovulation, AhR signaling, and oxidative stress.

| **Gene name** | **Description** | **Functions and references** |
| --- | --- | --- |
| *Pgr* | Progesterone receptor | A key LH/hCG-induced ovulatory gene that mediates progesterone receptor signaling in preovulatory follicles and is required for follicle rupture during ovulation (1). |
| *Ptgs2* | Prostaglandin-endoperoxide synthase 2 | LH/hCG-induced enzyme required for prostaglandin production, cumulus expansion, and follicle rupture during ovulation (2). |
| *Has* | Hyaluronan synthase 2 | LH/hCG-induced enzyme that drives hyaluronan production and cumulus expansion, a key process required for successful ovulation (3). |
| *Ereg* | Epiregulin | LH/hCG-induced EGF-like ligand that activates EGFR signaling to promote cumulus expansion, oocyte maturation, and ovulation (4). |
| *Sult1e1* | Sulfotransferase family 1E member 1 | hCG-induced estrogen sulfotransferase that supports cumulus expansion and ovulation (5). |
| *Star* | Steroidogenic acute regulatory protein | LH/hCG-induced steroidogenic regulator that promotes cholesterol transport and progesterone production during follicle luteinization (6). |
| *Ahrr* | Aryl hydrocarbon receptor repressor | AhR-inducible repressor that provides negative feedback to limit AhR signaling and xenobiotic/stress-responsive gene expression (7). |
| *Cyp1a1* | Cytochrome P450 family 1 subfamily A member 1 | Canonical AhR target gene involved in xenobiotic metabolism and is commonly used as a marker of AhR pathway activation (8) |
| *Cyp1b1* | Cytochrome P450 family 1 subfamily B member 1 | AhR-inducible cytochrome P450 enzyme involved in xenobiotic metabolism and oxidative stress-associated responses (9). |
| *Sod1* | Superoxide dismutase 1 | Cytosolic antioxidant enzyme that converts superoxide radicals into hydrogen peroxide, serving as a first-line defense against oxidative stress (10). |
| *Sod2* | Superoxide dismutase 2 | Mitochondrial antioxidant enzyme that detoxifies superoxide radicals and protects cells from mitochondrial oxidative stress (10). |

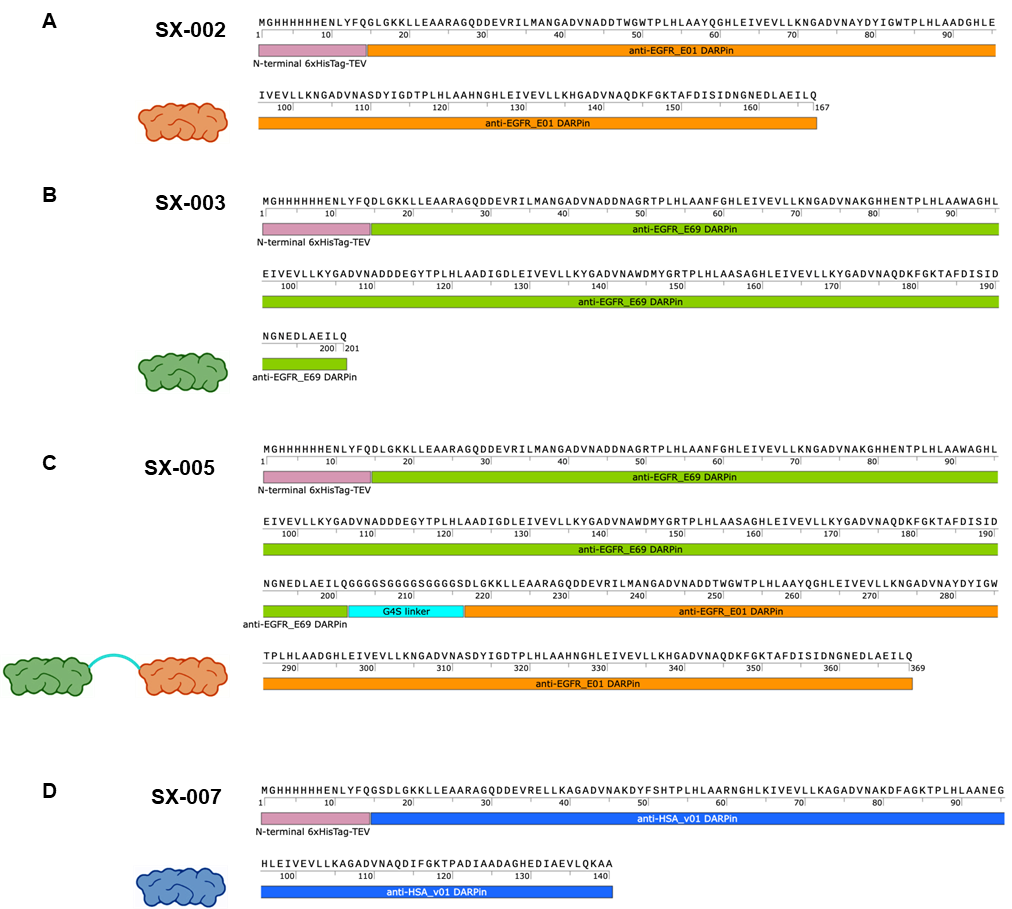

**Figure S1.** Annotated amino acid sequences of the expanded DARPin panel utilized in this study, detailing domain structures for (**A**) the monovalent SX-002, (**B**) the monovalent SX-003, (**C**) the flexible-linker bispecific SX-005, and (**D**) the anti-HSA negative control DARPin SX-007.

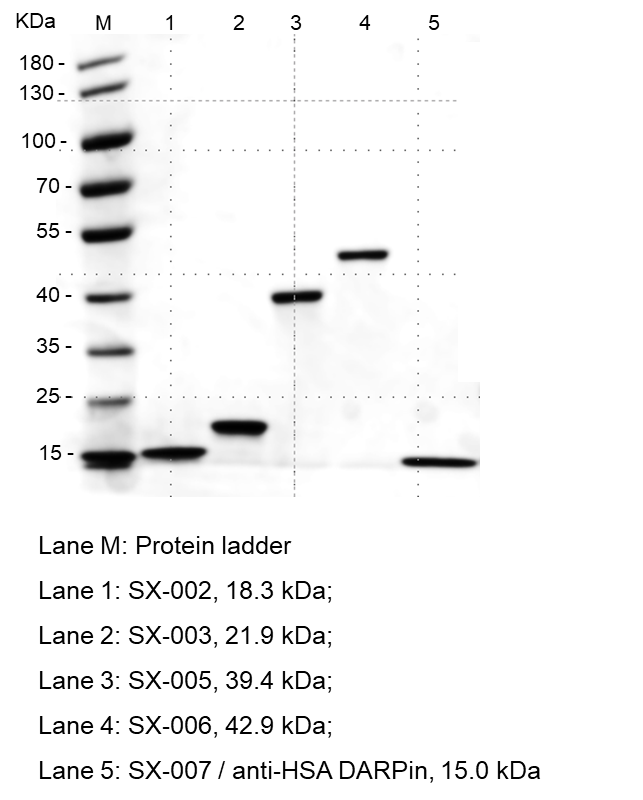

**Figure S2.** SDS-PAGE analysis of purified recombinant DARPins. Purified recombinant DARPin proteins were resolved by SDS-PAGE and visualized by Coomassie staining.

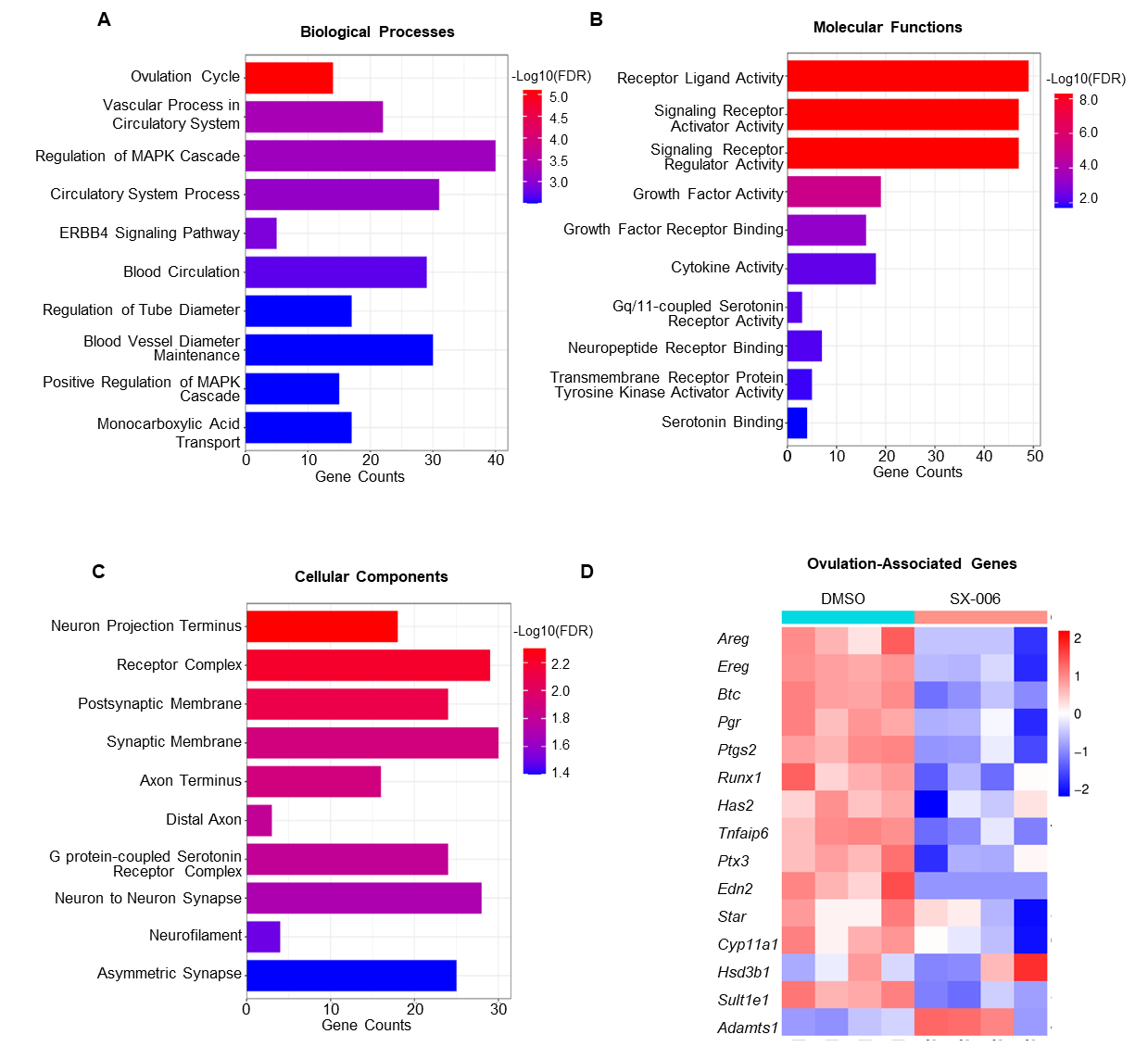

**Figure S3** GO enrichment analysis of SX-006–downregulated genes and ovulation-associated gene expression. GO enrichment analysis was performed using downregulated DEGs identified in SX-006-treated follicles at 4 h post-hCG. The top 10 enriched GO terms are shown for biological processes (A), molecular functions (B), and cellular components (C). The x-axes represent the number of downregulated DEGs associated with each GO term. Bar colors indicate enrichment significance expressed as −log10(FDR). (D) (D) Heatmap showing the expression patterns of major ovulatory marker genes identified by RNA-seq. Expression values were transformed to row Z-scores. Red and blue indicate relatively high and low expression levels, respectively.

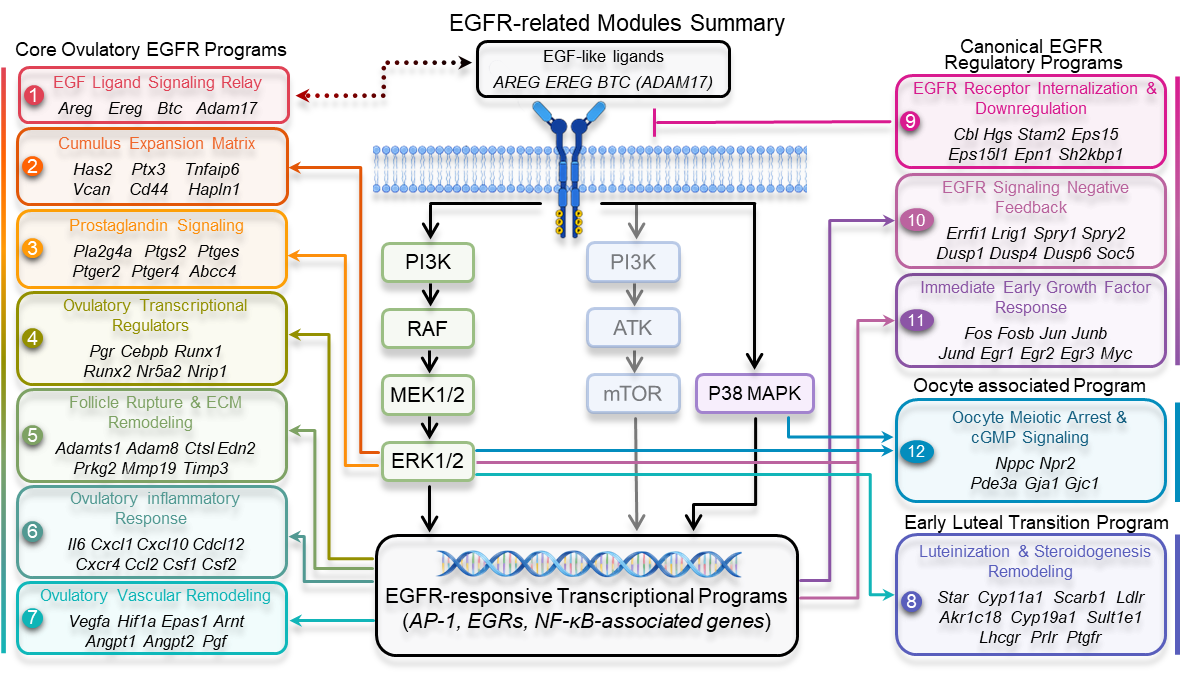

**Figure S4** Literature-curated EGFR-associated modules used for pathway-level analysis of ovulatory signaling. Schematic overview of the 12 literature-curated EGFR-associated modules used for gene set enrichment analysis (GSEA). Modules were organized into four biological categories: core ovulatory EGFR programs (Modules 1–7), early luteal transition program (Module 8), canonical EGFR regulatory programs (Modules 9–11), and oocyte associated program (Module 12). The diagram summarizes established relationships between LH-induced EGF-like ligands, EGFR downstream signaling pathways, EGFR-responsive transcriptional programs, and ovulation-associated biological processes based on previously published studies. Solid arrows indicate direct or well-established signaling relationships, whereas module connections represent literature-supported functional associations rather than direct molecular interactions. This framework was used to construct custom gene sets for downstream GSEA analysis.

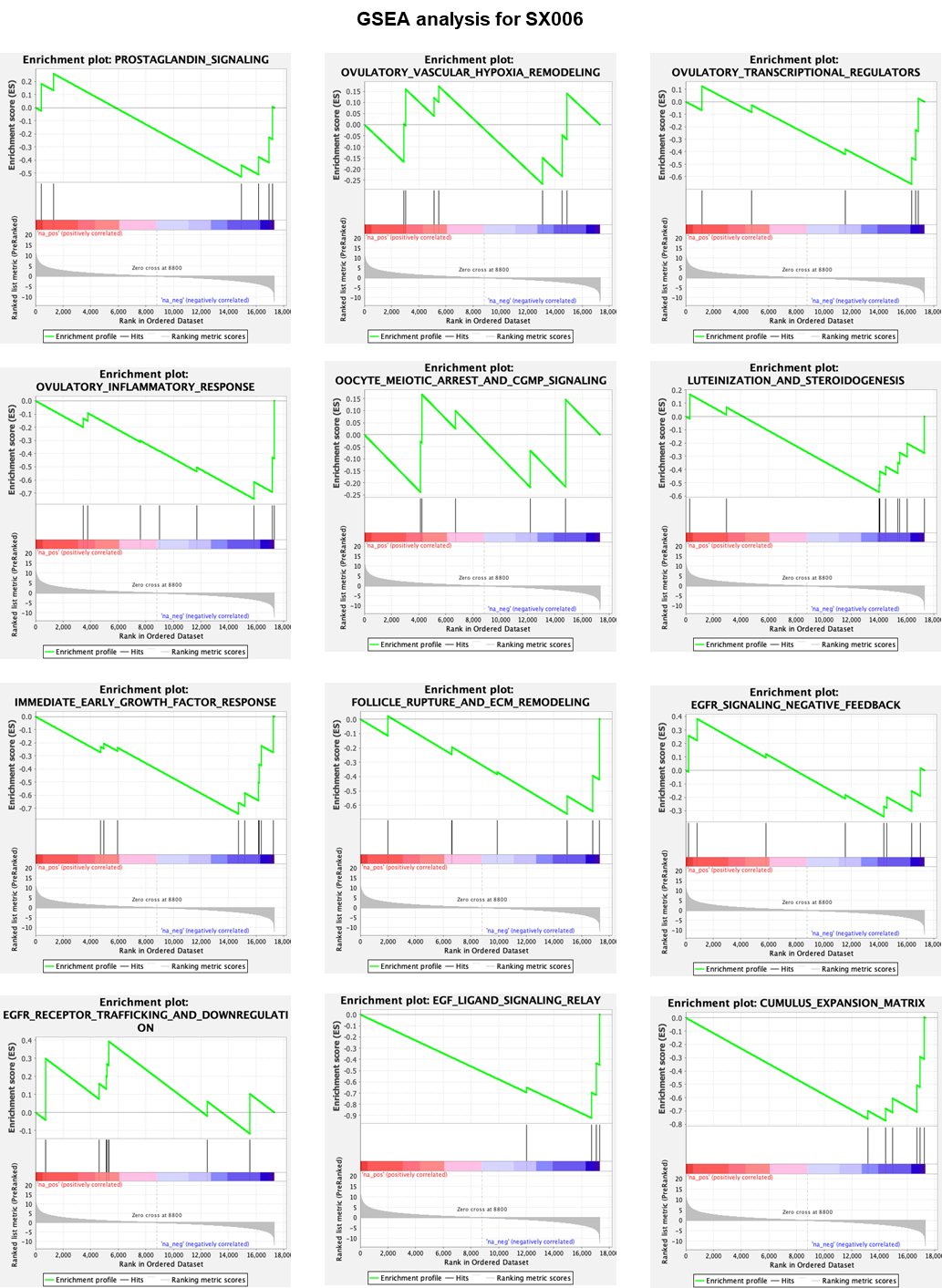

**Figure S5** GSEA enrichment plots of literature-curated EGFR-associated transcriptional modules in SX-006-treated follicles. Gene set enrichment analysis (GSEA) was performed using ranked RNA-seq data from SX-006-treated follicles compared with control follicles collected 4 h after hCG stimulation. The enrichment plots show representative EGFR-associated modules related to EGF ligand signaling relay, cumulus expansion matrix, prostaglandin signaling, ovulatory transcriptional regulators, follicle rupture and extracellular matrix remodeling, ovulatory inflammatory response, ovulatory vascular remodeling, oocyte meiotic arrest and cGMP signaling, luteinization and steroidogenesis remodeling, immediate-early growth factor response, EGFR signaling negative feedback, and EGFR receptor trafficking and downregulation. Vertical black lines indicate the positions of module genes within the ranked gene list. Positive and negative enrichment indicate relative enrichment at the top or bottom of the ranked dataset, respectively.

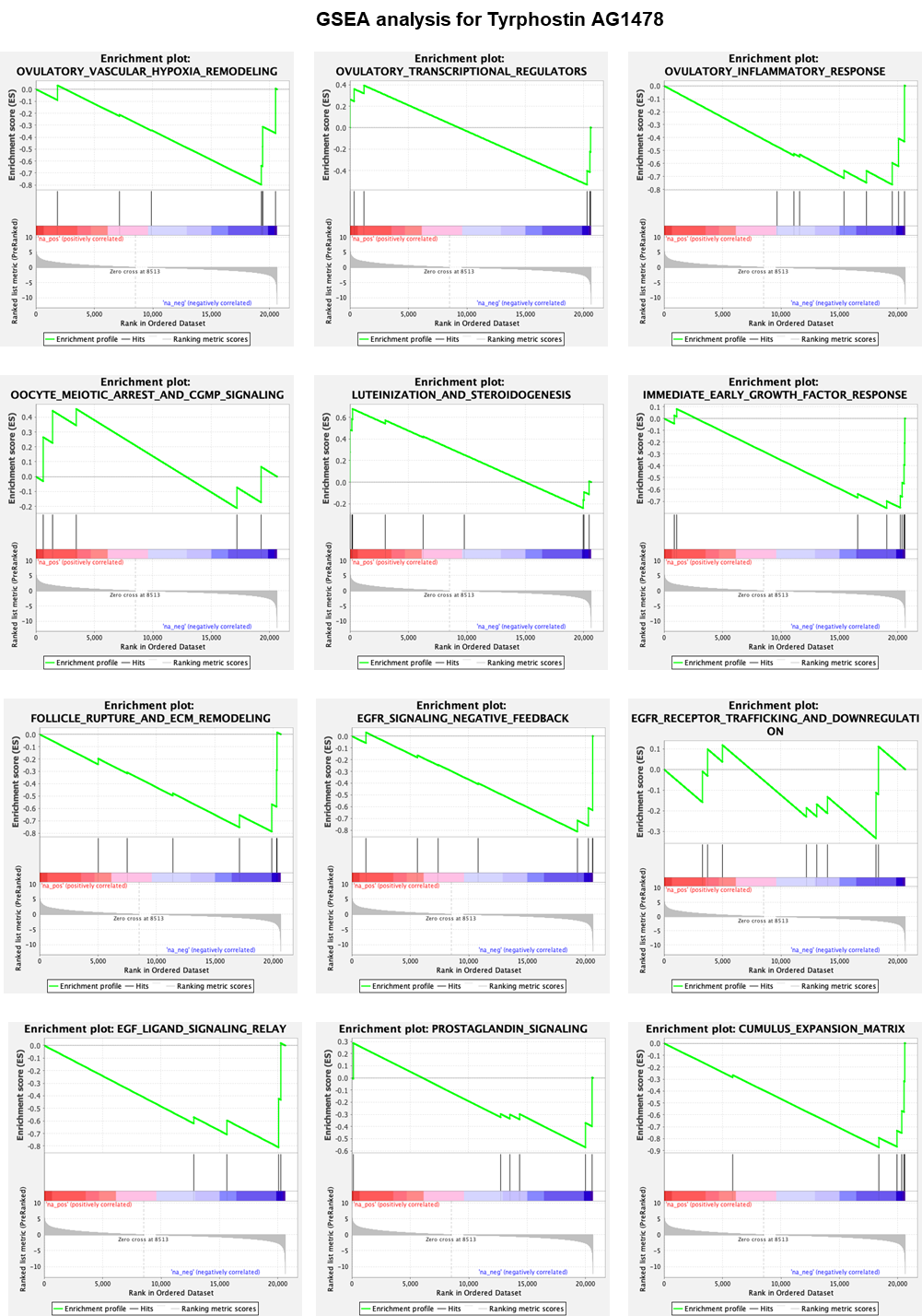

**Figure S6** GSEA enrichment plots of literature-curated EGFR-associated transcriptional modules in Tyrphostin AG1478-treated follicles. GSEA was performed using ranked RNA-seq data from Tyrphostin AG1478-treated follicles compared with control follicles collected 4 h after hCG stimulation. Vertical black lines indicate the positions of module genes within the ranked gene list.
